# In Silico Evaluation of Plant Nitrification Suppression Effects on Agroecosystem Nitrogen Loss

**DOI:** 10.1101/2022.02.28.482267

**Authors:** Melannie D. Hartman, Mark Burnham, William J. Parton, Adrien Finzi, Evan DeLucia, Wendy H. Yang

## Abstract

Nitrification regulates potential for nitrogen (N) loss from ecosystems because it converts ammonium to nitrate, which is susceptible to leaching and gaseous emissions. Crops can suppress the microbes that perform nitrification by exuding nitrification-inhibiting compounds from their roots and taking up available ammonium, the substrate for nitrification. However, the effect of nitrification suppression on agroecosystem N losses remains poorly characterized, and a lack of temporal synchrony between nitrification, N losses, and nitrification suppression by plants could limit the effect of nitrification suppression. We used the DayCent-CABBI model to evaluate the effectiveness of the suppression of nitrification by sorghum to reduce N_2_O emissions and nitrate leaching in an energy sorghum/soybean rotation at the Energy Farm in Urbana-Champaign, IL. We simulated nitrification suppression at the measured levels (measNS) and at the maximum measured level applied to the entire growing season (maxNS), and we also explored ways to better utilize nitrification suppression by altering the timing of UAN fertilizer applications. Model experiments showed that most nitrification occurred immediately after fertilizer was applied, whereas nitrification suppression begins to ramp up more than a month after planting. On an annual basis, measNS experiments showed a 1-2% reduction in annual N_2_O emissions relative to no nitrification suppression (noNS), and maxNS experiments showed a 4-9% reduction in annual N_2_O emissions relative to noNS. Both nitrification suppression levels showed < 1% reduction in nitrate leaching. Altering the timing of fertilizer applications to better synchronize nitrification suppression with high soil ammonium levels had mixed effects on annual N_2_O emissions and nitrate leaching and sometimes resulted in increased N losses. The timing of simulated N_2_O emissions shifted with the timing of fertilization, and N_2_O emissions from denitrification increased when N_2_O emissions from nitrification decreased. Increasing N retention during the non-growing season may be more effective and growing-season nitrification suppression for reducing annual N losses in the rainfed Midwest, particularly for NO_3_^-^ leaching in the early spring. Optimizing the timing of nitrification suppression alongside off-season N retention strategies would best improve the N sustainability of agroecosystems.

## Introduction

Controlling nitrification, the microbial process primarily responsible for soil nitrate (NO_3_^-^) production, is key to reducing nitrogen (N) losses from agroecosystems. Leaching of NO_3_^-^ from agricultural fields reduces available N to support crop yield and represents a significant environmental pollutant that contributes to downstream eutrophication (Van Meter et al. 2017). Nitrification also contributes to soil emission of nitrous oxide (N_2_O), a potent greenhouse gas and precursor to stratospheric ozone-depleting compounds, by directly producing N_2_O as a by-product and fueling denitrification through the production of its substrate, NO_3_^-^ (Farquharson 2016). Synthetic nitrification inhibitors can reduce nitrification rates and thus NO_3_^-^ leaching and N_2_O emissions from agricultural fields (Di and Cameron 2002b, Gilsanz et al. 2016) but the cost of the inhibitors relative to variable gains in yield has hindered their widespread use (Yang et al. 2016). In recent years, the suppression of rhizosphere nitrification by plants, including biological nitrification inhibition (BNI) by plant root exudates, has received considerable attention as a more cost effective alternative (Subbarao et al. 2013b, Coskun et al. 2017, Nardi et al. 2020). While BNI has been demonstrated in laboratory and greenhouse settings (Subbarao et al. 2013a, Janke et al. 2018), the effect of plant nitrification suppression on N_2_O emissions and NO_3_^-^ leaching has remained uncertain (Teutscherová et al. 2021, Burnham et al. 2022).

Plants can both indirectly and directly suppress nitrification in the rhizosphere, the zone of soil within approximately 3 mm of fine roots (Finzi et al. 2015). Plant roots indirectly suppress nitrification by creating conditions in the rhizosphere that reduce the activity of nitrifying bacteria and archaea. For example, plant roots and microbes compete for mineral nitrogen in the rooting zone, which can significantly reduce the availability of ammonium (NH_4_^+^) (Inselsbacher et al. 2010) and lead to lower nitrification rates in the rhizosphere (Stienstra et al. 1994, Booth et al. 2005). In addition, heterotrophic microbes can potentially outcompete autotrophic nitrifiers for NH_4_^+^ in the rhizosphere as a result of dead roots and root exudates contributing to high organic carbon availability that favors heterotrophic activity (Elrys et al. 2021). Plants can also exhibit BNI through the release of root exudates demonstrated to directly inhibit nitrifier activity in laboratory assays in which collected root exudates are applied to soil or pure cultures of the ammonia oxidizer, *Nitrosomonas europaea* (Subbarao et al. 2013a, Coskun et al. 2017, Janke et al. 2018). Sorghum (*Sorghum bicolor* (L.) Moench) is one of the best studied plants with BNI capability, directly inhibiting nitrifiers by exuding sorgoleone, sakuranetin and methyl 3-(4-hydroxyphenyl) propionate (MHPP) from its roots (Subbarao et al. 2013a). Other grass species, including wheat, rye, and wild grasses, also have the capacity for direct BNI, suggesting that the trait may be widespread (O’Sullivan et al. 2016, Coskun et al. 2017, Subbarao et al. 2021). These indirect and direct effects of plants on nitrification can be difficult to partition (Teutscherová et al. 2021), particularly in field studies (Burnham et al. 2022), so we consider them together as plant nitrification suppression.

Nitrification suppression is dynamic throughout the growing season over the lifespan of a plant (Nardi et al. 2020, Burnham et al. 2022), contributing to uncertainty about its effects on N losses from agricultural fields. Plant N demand changes based on growth stage (Moll et al. 1982, van Oosterom et al. 2010, Ciampitti and Vyn 2011), resulting in temporal variability in plant competition for NH_4_^+^ in the rhizosphere and indirect suppression of nitrification (Burnham et al. 2022). Furthermore, changes in the amount and composition of root exudates throughout annual crop development can lead to greater direct BNI later in the plant’s life cycle (Zakir et al. 2008, Chaparro et al. 2013, Sarr et al. 2019, Nardi et al. 2020). Finally, as root biomass of annual crops increases over the growing season (Black et al. 2017), increasing rhizosphere volume presumably leads to greater indirect and direct suppression of nitrification at the field scale as the growing season progresses. Together these factors suggest little nitrification suppression early in the growing season, with increasing nitrification suppression as the crops develop over the growing season.

The effectiveness of plant nitrification suppression in reducing agroecosystem N losses likely depends on the synchronization of temporal dynamics in nitrification rates and nitrification suppression over the growing season. In the agriculturally-intensive Midwestern US, the majority of fertilizer N (∼75%) is applied at or before the planting of annual crops (Cao et al. 2018), when plant N uptake is low or non-existent. This leaves microbes as the main consumers of soil NH_4_^+^ prior to crop establishment, with nitrification rates many-fold higher than plant uptake or microbial assimilation (Inselsbacher et al. 2010). Plant root biomass and BNI compound release are also lower early in the growing season (Zakir et al. 2008, Black et al. 2017, Sarr et al. 2019), so both direct and indirect mechanisms of nitrification suppression by plants are minimal at the time of fertilizer-induced stimulation of nitrification. In addition, spring and early-summer rainfall account for approximately one-third of total annual precipitation in this region (Huff and Angel 1992), with the intensity and frequency of early growing season rain events increasing over the past few decades (Dai et al. 2016). This combination of fertilizer application and wet soil conditions in the early growing season could minimize the potential for plant nitrification suppression to reduce NO_3_^-^ production and the resulting N_2_ O emissions and NO_3_^-^ leaching losses from intensively-managed annual cropping systems.

Here we use in silico experiments to test the hypothesis that greater synchronization between plant nitrification suppression and gross nitrification rates leads to greater reductions in agroecosystem N losses via soil N_2_O emissions and NO_3_^-^ leaching. We specifically evaluated the potential for nitrification suppression by energy sorghum to reduce field-scale nitrogen losses in the Midwest U.S. Energy sorghum, an emerging bioenergy feedstock crop, is a high-yielding hybrid of biomass and sweet sorghum optimized for high vegetative growth and stem juice production. We calibrated and validated the DayCent-CABBI process-based biogeochemical model for energy sorghum using field measurements from two energy sorghum experiments at the University of Illinois Energy Farm in 2018 and 2019, including estimates of plant nitrification suppression through the growing season (Moore et al. 2020b, Quinn 2021, Schetter et al. 2021, Burnham et al. 2022). To test how the timing of nitrification suppression affects N losses, we compared model experiments all simulating conventional fertilizer application at the time of planting but with different timing and magnitude of nitrification suppression (i.e., no nitrification suppression, with measured levels of nitrification suppression that changed over the growing season and varied inter-annually, and with a consistently high level of nitrification suppression starting one month after planting). To test how the timing of fertilizer-induced stimulation of nitrification affects N losses, we compared model experiments all simulating measured levels of nitrification suppression but with different timing of fertilizer application (i.e., conventional fertilizer application at the time of planting, with delayed fertilization by 3-weeks and by 5-weeks, and with split fertilizer application). In addition to evaluating annual N losses in the 2018 and 2019 sorghum years, we also examined how long-term weather variability affects nitrification suppression in a sorghum-soybean rotation through 2030.

## Methods

### Site description

Field data were collected at the University of Illinois Energy Farm in Urbana, IL, USA (N 40.063607, W 88.206926). The predominant soil type at this site is the very deep, poorly drained Drummer silty clay loam (fine-silty, mixed, superactive, mesic Typic Endoaquolls). At this location, the mean annual temperature (2010-2019) was 9.0° C and the mean annual precipitation was 1023 mm, with a mean growing season temperature and precipitation of 17.0° C and 598 mm, respectively (NOAA National Climatic Data Center station ID USC00118740, 3.6 km from the field site). Daily weather data (2008-2020) for model simulations were obtained from a weather station onsite at the Energy Farm. Mean annual N deposition was estimated at 8.5 kg N ha^-1^ y^-1^ based on 5.0 kg N ha^-1^ y^-1^ wet N deposition (NH_4_–N + NO_3_–N) from 2000-2019 obtained from the National Atmospheric Deposition Program for Bondville, Illinois, which is approximately 15 km from the study site (http://nadp2.slh.wisc.edu/data/sites/siteDetails.aspx?net=NTN&id=IL11, accessed December 4, 2020), and an assumption that dry deposition was 70% of wet deposition (McIsaac et al. 2002; USEPA, 2007).

### Field experiment data

Field experiment data on mineral nitrogen pools, nitrification suppression, and aboveground biomass for calibrating the model simulations were obtained from a two-year fertilization field trial of energy sorghum conducted in 2018 and 2019. Field trial descriptions and methods for sample collection and analyses are provided in detail by Schetter et al. (2021) and Burnham et al. (2022). In brief, because sorghum is grown in annual rotation with soybean, the two years of the field trial were established on two separate but adjacent fields (200 m apart) with similar soils but slightly different historical management (Table 1, Table 2). This allowed each year of the field trial to occur in sorghum plots preceded by soybean the previous growing season. Within each field, replicated plots (n = 4) were set up in a randomized block design with four different levels of urea-ammonium-nitrate (1:1:1) (UAN) fertilizer that was applied within a week of planting: 0, 56, 112, and 168 kg N ha^-1^ y^-1^. These trials also included four energy sorghum genotypes, TX17500, TX17600, TX17800, and TX08001. However, we did not detect any differences in nitrification suppression between genotypes (Burnham et al. 2022), and all genotypes had similar growth curves and biomass yields (Schetter et al. 2021); thus, we averaged our data across the four genotypes. To evaluate changes in soil mineral nitrogen pools after the growing season, we measured potassium chloride (KCl) extractable NO_3_^-^ and NH_4_^+^ in the top 30 cm of soil prior to fertilization and from 0-10 cm, 10-30 cm, and 30-50 cm depth after sorghum harvest. Nitrification suppression measurements occurred three times each growing season, the first occurring about a month after sorghum was planted (late June 2018, early July 2019), the second in the middle of the growing season (late July 2018, early August 2019), and the last near the end of the growing season (late August 2018, early September 2019). We performed *ex situ* potential nitrification assays on bulk and rhizosphere soils using a method adapted from Belser and Mays (1980), estimating nitrification suppression rates as the difference between rhizosphere and bulk soil potential nitrification rates (Burnham et al. 2022).

**Table 1.** Summary of model nitrification suppression and fertilizer timing experiments. These treatments applied to years that sorghum was grown starting in 2018 or 2019.

| Primary experiment<br>(abbreviation) | Sorghum UAN fertilizer application rates<br>and timing for the experiment | Nitrification suppression level |
| --- | --- | --- |
| Measured nitrification suppression<br>(measNS) | 0, 56, 112, 168<br>kg N ha<sup>-1</sup> y<sup>-1</sup> within a few days of planting<br>sorghum | nitrification suppression as was measured<br>in the field |
| No nitrification suppression<br>(noNS) | 0, 56, 112, 168<br>kg N ha<sup>-1</sup> y<sup>-1</sup> within a few days of planting<br>sorghum | no reduction in gross nitrification |
| Maximum nitrification suppression<br>(maxNS) | 0, 56, 112, 168<br>kg N ha<sup>-1</sup> y<sup>-1</sup> within a few days of planting<br>sorghum | 70% reduction in gross nitrification<br>starting 1 month after planting and<br>continuing 16 weeks, to the end of the<br>growing season |
| Delay one-time fertilizer application 3<br>weeks (measNS_dly3wk) | 56, 112, 168<br>kg N ha<sup>-1</sup> y<sup>-1</sup> three weeks after actual<br>fertilization date | nitrification suppression as was measured<br>in the field |
| Delay one-time fertilizer application 5<br>weeks (measNS_dly5wk) | 56, 112, 168<br>kg N ha<sup>-1</sup> y<sup>-1</sup> five weeks after actual<br>fertilizer date | nitrification suppression as was measured<br>in the field |
| Split fertilizer application (measNS_split) | 56, 112, 168<br>kg N ha<sup>-1</sup> y<sup>-1</sup> where half is applied a few<br>days of planting sorghum, the other half 4<br>weeks later | nitrification suppression as was measured<br>in the field |

**Table 2.**
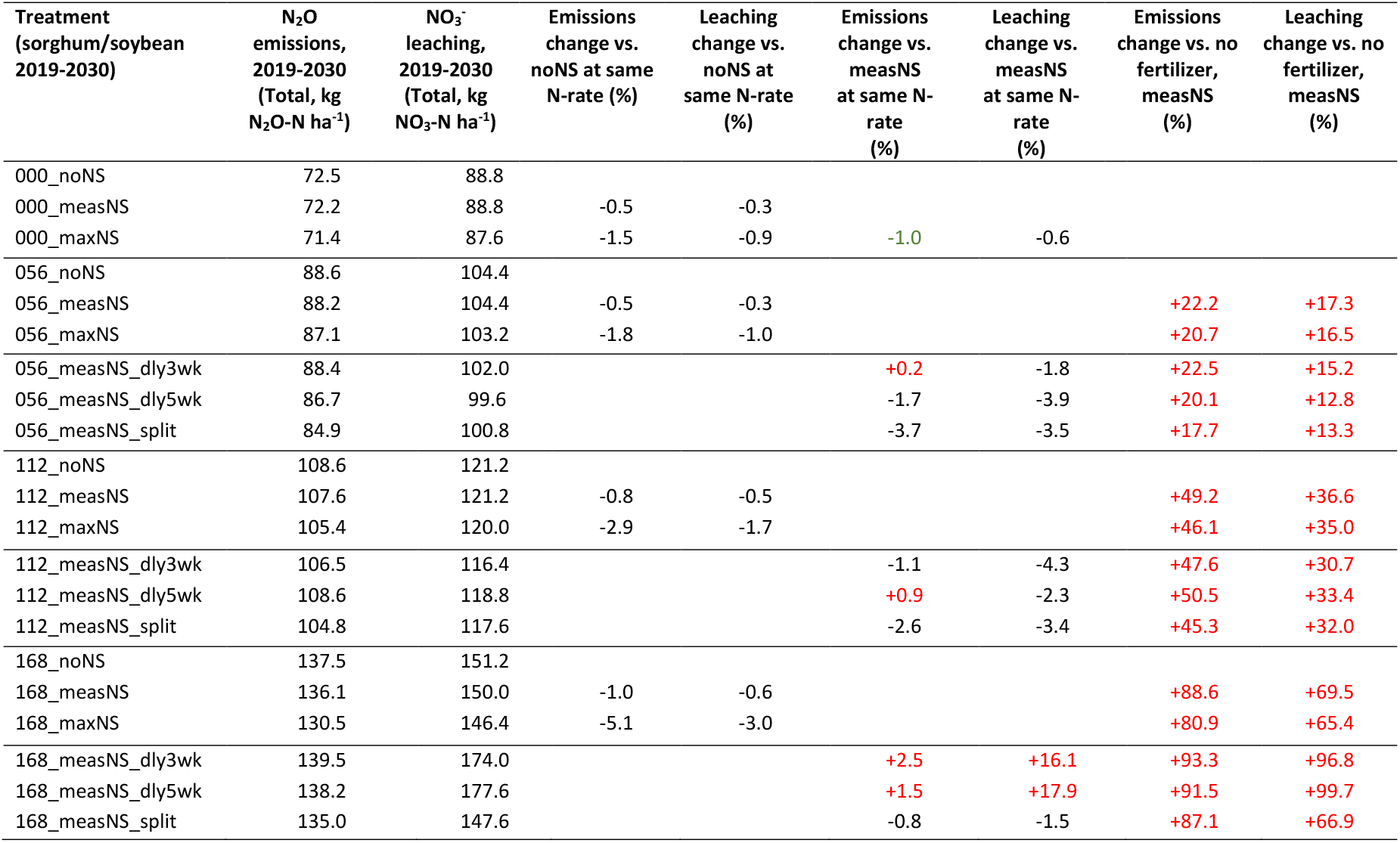
Simulated cumulative N_2_O emissions and nitrate leaching from 2019-2030, where sorghum was grown in odd years and soybean was grown in even years. Experiments include sorghum nitrification suppression (noNS = no nitrification suppression, measNS = measured nitrification suppression (2019), maxNS = maximum nitrification suppression) and fertilizer timing experiments (dly3wk = 3-week delay, dly5wk=5 week delay, split = split fertilizer application) at four UAN fertilizer application amounts (N-rate = 0, 56, 112, or 168 kg N ha^-1^ y^-1^). No nitrification suppression or fertilizer was applied in soybean years.

Field experiment data on soil N_2_O emission and root biomass for calibrating the model simulations were obtained from five sorghum plots located within 200-1000 m of the fertilization field trial and on the same soil series as the field trial. These plots include one 8.7 ha plot and four replicated 1.7 ha plots planted with the TX17500 genotype, which was also planted within the sorghum field trial, and fertilized with UAN at 112 kg N ha^-1^ y^-1^ rate at planting, similar to the field trial. We measured N_2_O fluxes using the static flux chamber method 12 times throughout the 2018 growing season (May-October), and cumulative growing season N_2_O emissions were calculated by linearly interpolating daily N_2_O flux rates between sample dates. Details about these measurements and plots are provided by Moore et al. (2020b). Root biomass was measured at 0-10 cm, 10-20 cm, 20-30 cm, 30-50 cm, and 50-100 cm depth increments in soil cores collected from 2 locations near the center of each plot, with one core taken from an alley and one taken from a planted row. Fine (<2 mm) and coarse root biomass was separately quantified after manually extracting roots from soil cores and drying at 50°C for 72 hr (Quinn 2021).

### DayCent-CABBI model calibration and validation

We used DayCent, a daily time-step, process based biogeochemical model (Parton et al. 1998) that has been extensively calibrated and validated to simulate the effects of agricultural management on crop growth, long-term soil organic carbon dynamics (Six et al. 2004, Ogle et al. 2010) and N trace gas emissions (Del Grosso et al. 2005, Del Grosso et al. 2010). It is widely applied for quantification of greenhouse emissions budgets of varying production systems (Del Grosso et al. 2002, EPA 2018). The model was recently applied by to assess the sustainability of forage sorghum in the semiarid southwest (Duval et al. 2015) and to investigate energy sorghum production emissions relative to corn at 3,265 sites across the rainfed United States (Kent et al. 2020). For the model experiments in this research, we used a new version of DayCent that was developed by the Center for Advanced Bioenergy and Bioproducts Innovation (DayCent-CABBI) based on the DayCent-Photo model version (Straube et al. 2018) and applied to simulate switchgrass and maize at the Energy Farm by Moore et al. (2020a).

The model was set up to simulate the historic management of the site (Table S1, S2). Sorghum 2018 and sorghum 2019 were on separate plots with somewhat different cropping history from 2011 - 2017, each sorghum crop was preceded by and followed the next year by a soybean crop (Table S1, S2). Correctly simulating crop yields, root biomass, and C:N ratios of plant material is necessary to accurately estimate plant N-uptake from soils and plant residue inputs to soils. The model was calibrated using measured sorghum yields, C:N of sorghum yields, measured root biomass, C:N of root biomass, cumulative N_2_O emissions during the 2018 growing season, and pre- and post-growing season ammonium and nitrate levels in the soil. To validate the model, parameters determined from the calibration were applied using all available 2019 data, including sorghum biomass yields, C:N of sorghum yields, and pre-growing season ammonium and nitrate levels. The model assumed a maximum rooting depth of 60 cm for both sorghum and soybeans with the majority of roots between 5 and 30 cm (Benjamin and Nielsen 2004, Quinn 2021). The distribution of roots affects both plant N uptake and transpiration. The simulated values used for calibration and/or validation were those from the measNS model experiments (measured nitrification suppression levels with no fertilizer delay, described below) since those experiments most closely replicated management that occurred in the field experiments.

### Model Experiments

We performed six primary model experiments to quantify the effect of nitrification suppression on N_2_O emissions (from both nitrification and denitrification), NO_3_^-^ leaching (nitrate exiting the soil profile), and sorghum aboveground net primary productivity (ANPP). In each primary experiment, we ran simulations for each sorghum growing season (2018 and 2019) with four levels of fertilizer applications (N-rates; 0, 56, 112, and 168 kg N ha^-1^ y^-1^) for each primary experiment (Table 3). To run the model experiments over a wider range of weather conditions, the model experiments were continued to the year 2030, starting with 2019 sorghum and alternating sorghum and soybean so that sorghum occurs on the odd calendar years and soybeans on the even calendar years. Weather from 2009-2018 was used for the 2021-2030 scenarios. Nitrification suppression levels during sorghum years were equivalent to those measured in sorghum in 2019, and there was no nitrification suppression or fertilization during soybean years.

The first set of three primary model experiments focused on different scenarios of the magnitude and timing of nitrification suppression: no nitrification suppression (noNS), measured nitrification suppression (measNS), and maximum nitrification suppression (maxNS). The noNS experiment represents actual crop management but assumes that no nitrification suppression occurs. This experiment serves as a baseline against which nitrification suppression effects on crop yield and agroecosystem N losses are assessed. The measNS experiment simulates the actual crop management and nitrification suppression levels that were measured in the field by Burnham et al. (2022) and begin approximately one month after planting sorghum (Table S2). The purpose of this model experiment is to evaluate reductions in N losses by measured levels of nitrification suppression, which peaks in the mid-growing season. The maxNS experiment simulates actual crop management but assumes a consistently high nitrification suppression level, a 70% reduction in nitrification that begins one month after planting sorghum (same date as the measNS experiment) and continues to the end of the growing season in mid-October (18 weeks). The purpose of this experiment is to evaluate the maximum possible benefit of BNI if it were able to achieve its highest measured level for the entire growing season. The effects of nitrification suppression (measNS and maxNS) are reported relative to no nitrification suppression (noNS) experiment results.

The second set of three primary model experiments focused on evaluating the potential for fertilizer management (timing) to maximize N loss reduction benefits from measured levels of nitrification suppression. We investigated three scenarios of delayed or split fertilizer application aimed at better synchronizing the fertilizer-induced stimulation of nitrification with the onset of nitrification suppression, which starts approximately 1 month after planting: 3-week delayed, 5-week delayed, and split fertilizer application. The 3-week delayed experiment simulated actual crop management except that the UAN fertilizer application occurred three weeks later than it was actually applied in the field experiments. The 5-week delayed experiment simulated actual crop management except that the UAN fertilizer application occurred five weeks later than it was actually applied in the field experiments. The split application experiment simulated actual crop management except that half the UAN fertilizer was applied on the date it was actually applied in the field, and the other half was applied four weeks later. The effects of altered fertilizer management are reported relative to the measNS experiment which differs from these three experiments only in the timing of fertilizer application.

## Results

### Model calibration and validation

The means of measured values of harvested sorghum biomass increased with N-rate, but these means were not significantly different from one another in either 2018 or 2019 (Figure S2 A, B). Simulated sorghum yield also increased with N-rate, and simulated values (one value per fertilization level per year) were not significantly different from mean observed values except at no fertilizer in 2019 (Figure S2 A, B). The C:N ratio of measured and simulated sorghum harvested biomass (2018) decreased with N-rate, and simulated values were within one standard error mean of measured values (Figure S2 C). The peak simulated root biomass for both N-rates (0.0 and 168 kg N ha^-1^ y^-1^) in 2018 were also within one standard error of the measured means (Figure S1 A), as was the simulated C:N ratios of root biomass (Figure S1 B).

Cumulative measured N_2_O emissions began May 5, 2018 (Time = 2018.34) for the 112 kg N ha^-1^ y^-1^ fertilizer treatment; therefore, simulated cumulative N_2_O emissions for sorghum 2018 from the measNS model experiment at the same N-rate that began on this same date were compared to the measured values (Figure S3). Though simulated cumulative emissions initially increased at a greater rate than measured emissions, simulated emissions are within the lower and upper bound of measured emissions by the beginning of June (Time = 2018.42). By the end of the growing season in mid-October (Time = 2018.8), simulated cumulative N_2_O emissions are comparable to the mean of measured emissions.

Pre-growing season measured ammonium and nitrate levels were made for the sorghum plots on May 18, 2018 and June 7, 2019, while post-growing season measurements were made on October 1, 2018 (Table S3). The depths over which measurements occurred varied and were not always directly comparable to DayCent results because DayCent simulates a single soil ammonium pool assumed to occur in the top 20 cm of soil while simulated nitrate pools exist for each of the 13 soil layers and extend through the soil profile down to 180 cm. Pre-growing season measured ammonium levels in the 0-15cm layer in 2018 (0.8 ± 0.3 kg N ha^-1^) and in the 0-30cm layer in 2019 (2.2 ± 0.2 kg N ha^-1^) were less than DayCent’s estimates of total soil ammonium (2.9 and 3.6 kg N ha^-1^ in 2018 and 2019, respectively) (Table S3). Pre-growing season measured nitrate levels in the 0-15 cm layer in 2018 (14.6 ± 1.3 kg N ha^-1^) were greater than DayCent’s estimate for the same depth (7.4 kg N ha^-1^). Pre-growing season measured nitrate levels in the 0-30cm layer in 2019 (25.6 ± 0.7 kg N ha^-1^) were close to DayCent’s estimate over the same depth (23.0 kg N ha^-1^). DayCent showed greater rooting zone depletion of soil ammonium and nitrate by the end of the growing season than measurements indicated. The 2018 post-growing season measurements of ammonium (25 ± 1.7 kg N ha^-1^) and nitrate (25 ± 7.6 kg N ha^-1^) in the top 50 cm of soil were greater than DayCent’s estimates, 2.0 kg N ha^-1^ for total ammonium and 1.2 kg N ha^-1^ (0-45 cm) for nitrate, however DayCent showed high nitrate levels deeper in the soil profile where measurements were not taken (Table S3). On average across all simulations, DayCent showed that approximately 35% of all soil nitrate existed in depths from 105 - 180 cm.

### Nitrification suppression effects on sorghum nitrogen losses in 2018 and 2019

On an annual basis, the measNS and maxNS experiments showed little to no reductions in gross nitrification rates, N losses, and ANPP relative to the noNS experiments across all four fertilizer application rates. In 2018 and 2019, measNS reduced gross nitrification rates for sorghum by 1% and 2-3%, respectively. In contrast, maxNS led to about 10 times greater effects on nitrification, reducing gross rates for sorghum by 11-14% and 7-12% in 2018 and 2019, respectively (Figure 1 A, B; Figure 2 A, B). Patterns in annual N_2_O emissions mirrored those in annual gross nitrification rates, with the measNS experiment showing a 1-2% reduction and the maxNS experiment showing a 4-10% reduction, with greater reductions as fertilizer application rates increased (Figure 1 C, D; Figure 2 C, D; Table S4). However, sorghum NO_3_^-^ leaching did not respond to nitrification level suppression in 2018 as NO_3_^-^leaching was limited to ∼2.7 kg N ha^-1^ due to relatively dry conditions in the mid-growing season (Figure 1 E; Figure 2 E). However, even in 2019 when more typical rainfall amounts led to NO_3_^-^ leaching of ∼16-17 kg N ha^-1^, both measNS and maxNS showed less than 0.5% reduction in NO_3_^-^ leaching (Figure 1 F; Figure 2 F, Table S4). In both 2018 and 2019, simulated ANPP from the noNS, measNS, and maxNS experiments were comparable for a given fertilizer level (Figure S4).

**Figure 1.**
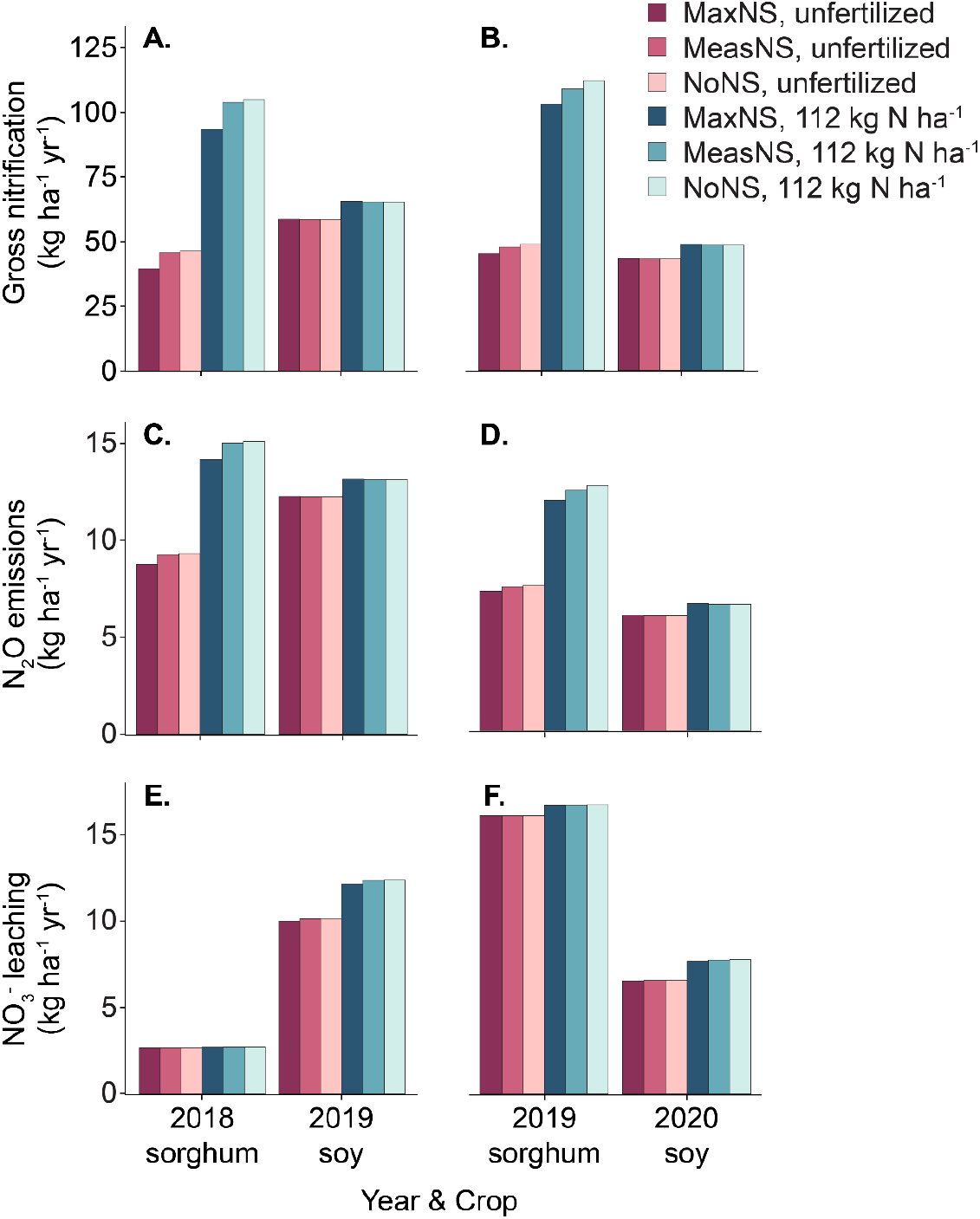
Simulated annual rates of gross nitrification (a, b), N_2_O emissions (c, d), and NO_3_^-^ leaching (e, f) under maximum (maxNS), measured (measNS), and no (noNS) suppression of nitrification by sorghum at 0 and 112 kg N ha^-1^ fertilization rates in 2018 (A, C, E) and 2019 (B, D, F) sorghum field trials. All rates were simulated during the sorghum growing season (2018 & 2019) and the following growing season when each trial was planted in soybean.

**Figure 2.**
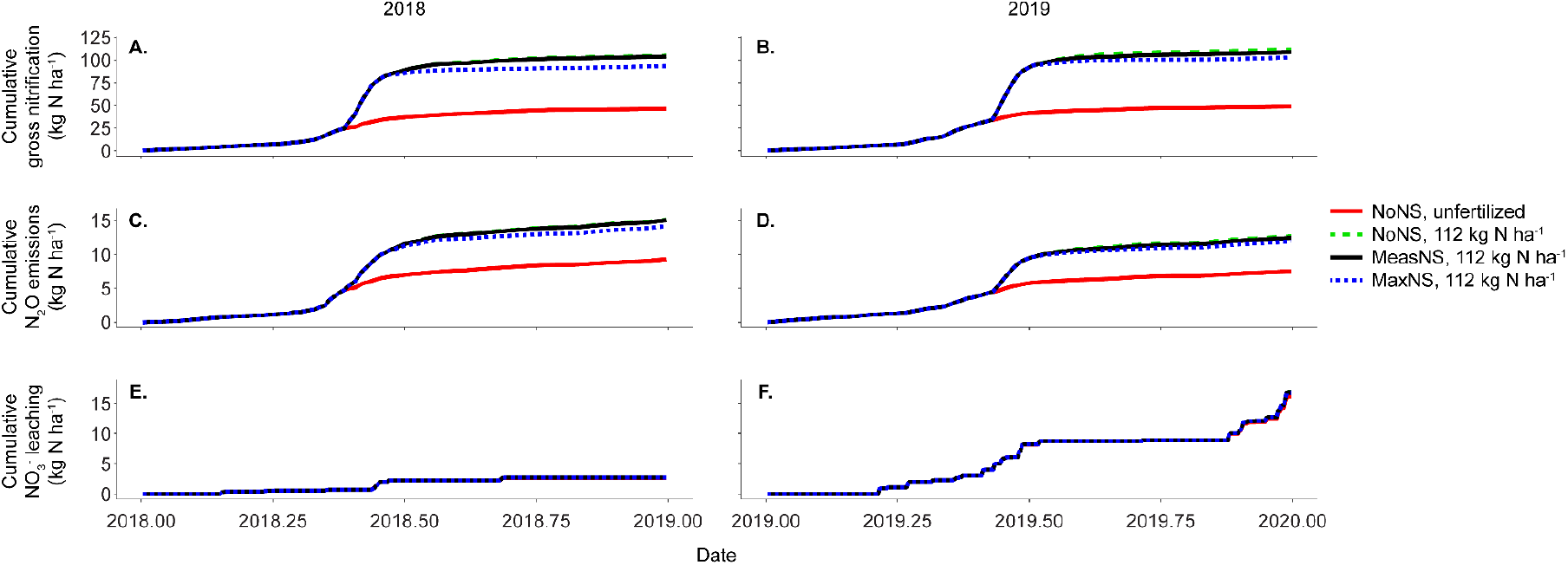
Simulated cumulative gross nitrification (A, B), N_2_O emissions (C, D), and NO_3_^-^ leaching (E, F) throughout the 2018 (A, C, E) and 2019 (B, D, F) growing seasons under maximum (maxNS), measured (measNS), and no (noNS) nitrification suppression by sorghum. Simulated rates are shown for noNS in unfertilized plots (red line) and for all simulated nitrification suppression rates in plots fertilized at a rate of 112 kg N ha^-1^.

Most nitrification occurred prior to establishment of annual sorghum crops and suppression of nitrification in both 2018 and 2019, regardless of fertilization rate (Figure 3). In fertilized plots, fertilizer was applied within a week of planting sorghum (Table S2) and nitrification increased dramatically immediately after fertilization in both in sorghum 2018 and 2019 (Figure 3 B). For the measNS model experiment with a fertilizer application rate of 112 kg N ha^-1^ y^-1^, 83% of annual gross nitrification had occurred by the time nitrification suppression was first measured on June 27^th^, and in 2019 88% of annual gross nitrification had occurred by the time nitrification suppression was first measured on July 8^th^ (Figure 3B). In both fertilized and unfertilized plots, little nitrification occurred after mid-season (Figure 3). Despite very low levels of nitrification after the early-season, nitrification was lower in the maxNS experiment than either noNS or measNS throughout the growing season, and this difference was greater in fertilized plots than unfertilized plots (Figure 3). Likewise, most cumulative nitrification, N_2_O emissions, and NO_3_^-^ leaching occurred early in the season in 2018 and 2019, and only the maxNS experiment maintained lower cumulative totals throughout the remainder of the season (Figure 2).

**Figure 3.**
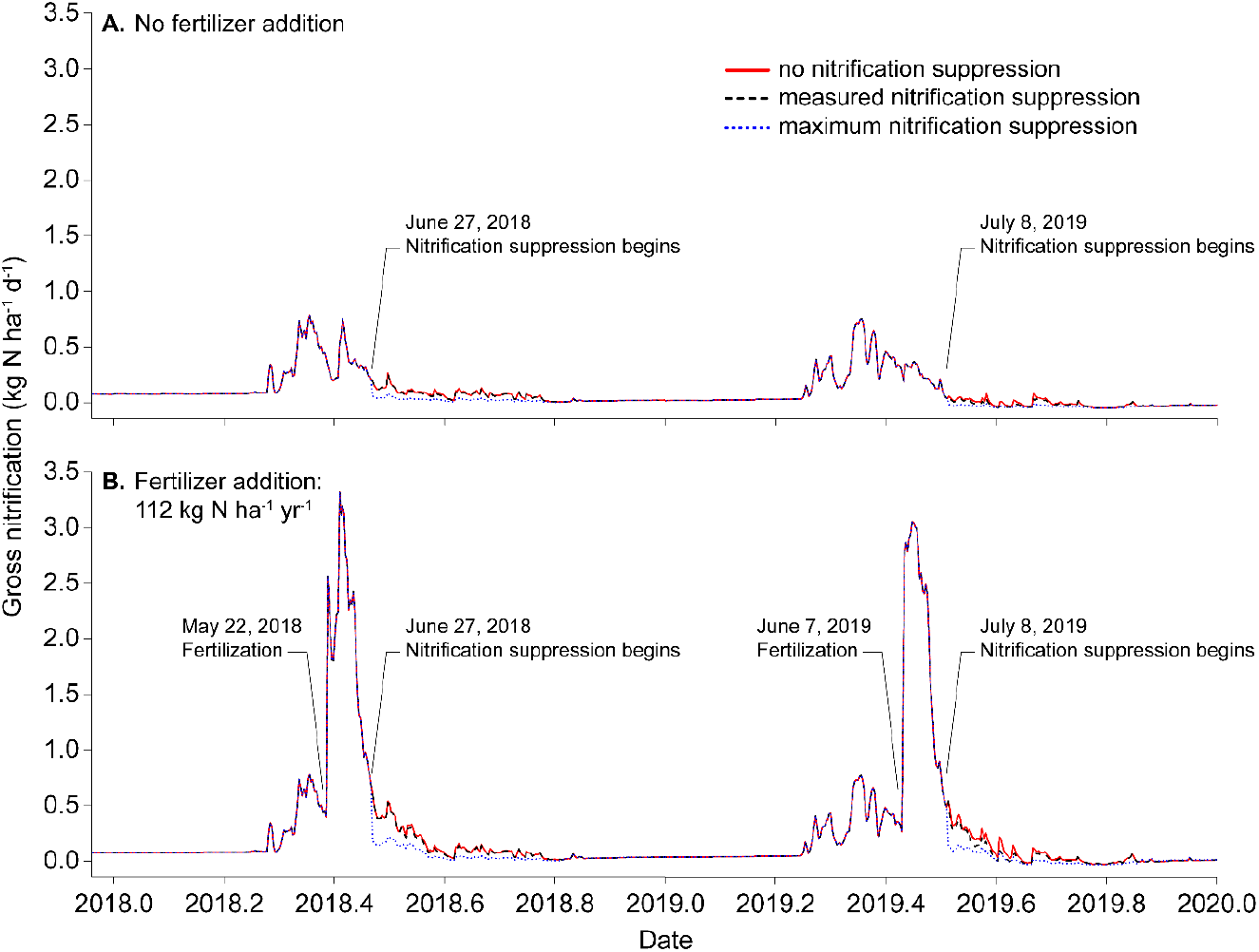
Simulated daily gross nitrification rate (kg N ha^-1^ d^-1^) during the 2018 and 2019 growing seasons in unfertilized (A) and fertilized (112 kg N ha^-1^, B) plots at three levels of nitrification suppression.

### Fertilizer timing effects on sorghum nitrogen losses in 2018 and 2019

Delaying fertilizer application affected simulated above-ground net primary production (ANPP) of sorghum in both years but splitting fertilizer applications had minimal effects on simulated ANPP compared to one-time fertilizer applications at a given total fertilizer application rate (Figure S4 C, D). In 2018 and 2019, delayed fertilizer applications increased ANPP by 3 - 6% for N-rates of 56 and 112 kg N ha^-1^ y^-1^, but decreased ANPP by as much as 7% for N-rates of 168 kg N ha^-1^ y^-1^, while splitting fertilizer applications increased ANPP from 0.0 to 2% relative to conventional fertilizer application around the time of planting (Figure S4 C, D).

In the fertilizer timing experiments, simulated gross nitrification increased immediately when UAN fertilizer was applied, both in sorghum 2018 and 2019 (Figure 4). Subsequently, the timing of fertilizer-induced gross nitrification was shifted so that it occurred when measured nitrification suppression commenced (Figure 4). However, delaying fertilizer application by three or five weeks increased annual gross nitrification rates 4.3-7.0% for sorghum 2018 relative to no delay in fertilizer application at all (Figure 5 A; Table S5). This led to a 3 – 9% increase in annual N_2_O emissions but only small changes (≤ 1%) in NO_3_^-^ leaching during 2018 (Figure 5 C, E; Table S5). In contrast, in 2019 when measured nitrification suppression levels were higher, delayed fertilizer application consistently decreased simulated gross nitrification rates by 1– 10% (Figure 4 B; Figure 5 B; Table S5). Additionally, the decrease in gross nitrification rates with delayed fertilizer addition in 2019 was greater at higher fertilization rates (Table S5). When 56 kg N ha^-1^ fertilization was delayed 3-weeks and 5-weeks, annual gross nitrification was 0.3% higher and 2.7% lower than without fertilization delay, respectively. However, when 168 kg N ha^-1^ fertilization was delayed 3- and 5-weeks, annual gross nitrification was 6.5% and 10.2% lower than without fertilization delay, respectively. At the same time, altering fertilization timing had variable effects on annual N_2_O emissions, which decreased 0.3-3.2% when 56 kg N ha^-1^ fertilization application was delayed but increased 1.0-1.5% when 112 or 168 kg N ha^-1^ application was delayed (Table S5). Delayed fertilizer application reduced NO_3_^-^ leaching by 0.5–3%, again with greater decreases in leaching with longer delays in application and higher application rates relative to fertilization at planting.

**Figure 4.**
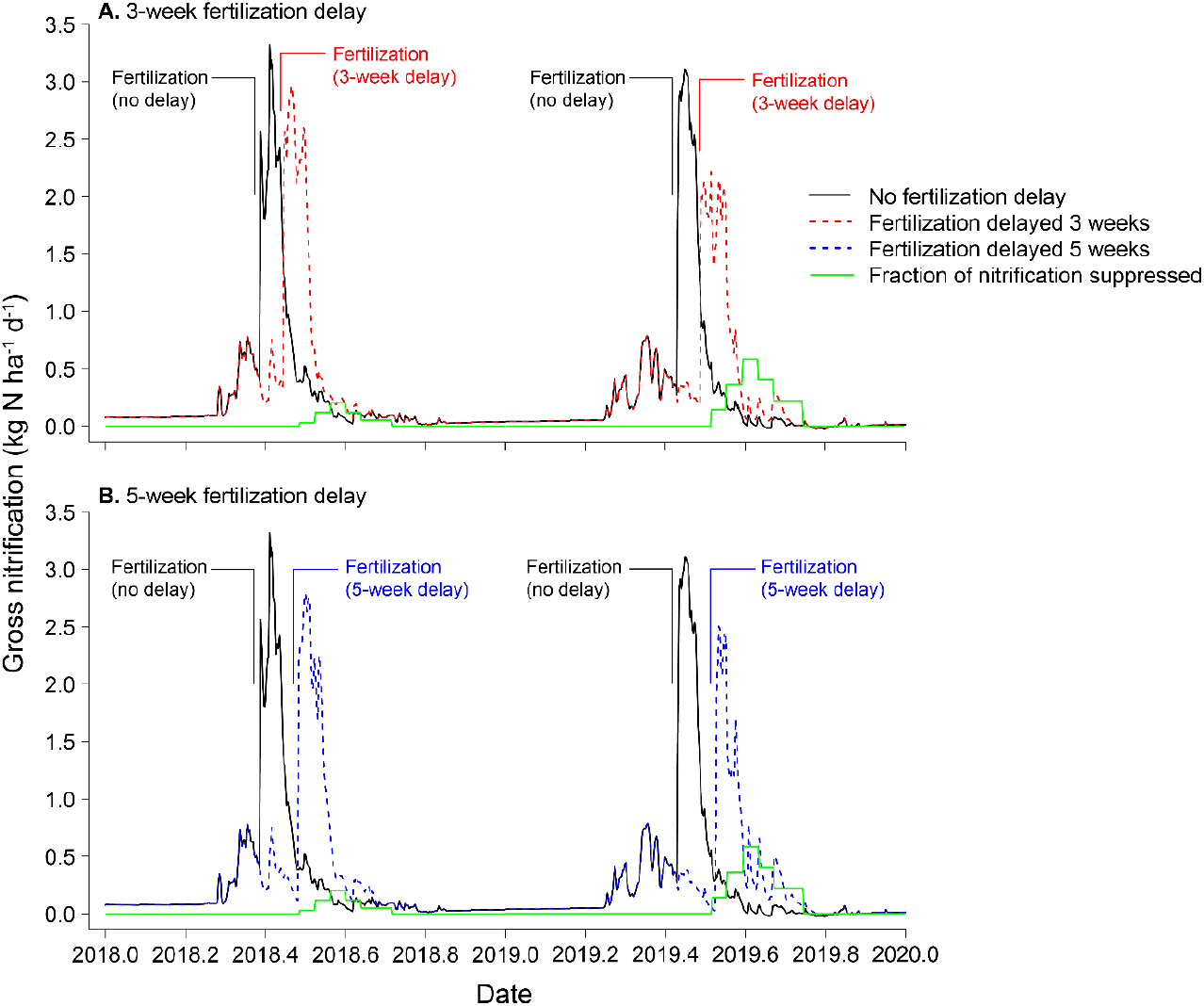
Simulated daily gross nitrification rate (kg N ha^-1^ d^-1^) under measured nitrification suppression (measNS) during the 2018 and 2019 growing seasons with fertilizer addition (112 kg N ha^-1^) delayed either 3 (A) or 5 (B) weeks. Solid black lines show gross nitrification rates without fertilizer delay, dashed lines show gross nitrification rates with delayed fertilizer addition, and the solid green lines show the timing and level of simulated nitrification suppression (measNS).

**Figure 5.**
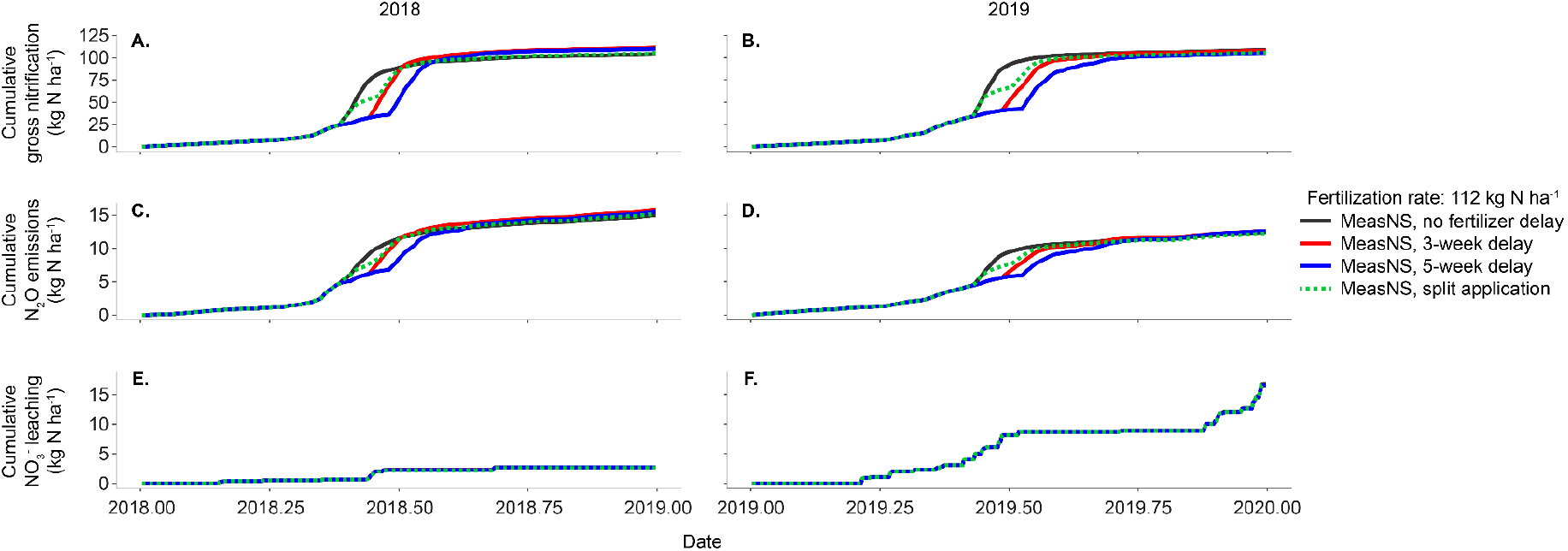
Simulated cumulative gross nitrification, N_2_O emissions, and NO_3_^-^ leaching throughout the 2018 and 2019 growing seasons with 112 kg N ha^-1^ fertilizer addition delayed 3 weeks or 5 weeks, or split into two additions, under measured (measNS) nitrification suppression by sorghum.

How split fertilizer application affected N losses from sorghum also differed between 2018 and 2019. In 2018, split application of 56 kg N ha^-1^ led to a 2.0% decrease in gross nitrification rate relative to fertilization at planting, but split application of higher amounts of fertilizer increased gross nitrification 0.8-0.9 %, and a similar pattern was observed in the effect of split fertilizer application on N_2_O emissions (Table S5). Also in 2018, split fertilizer application led to consistent 0.5-1.1% decreases in NO_3_^-^ relative to fertilization at planting, with greater decreases at higher fertilization rates. In 2019, split fertilizer application consistently reduced annual gross nitrification, annual N_2_O emissions, and NO_3_^-^ leaching rates. by 2-7%, annual N_2_O emissions by 1– 4%, and NO_3_^-^ leaching by < 1% (Table S5). Split fertilizer application of 56 kg N ha^-1^ reduced gross nitrification 7.0% and N_2_O emissions 4.0% compared to fertilization at planting, but these reductions declined to 1.6-2.7% for nitrification and 0.8-1.6% for N_2_O emissions with higher fertilizer application rates (Table S5). In contrast, 2019 NO_3_^-^ leaching was consistently reduced by 0.4-0.6% by splitting fertilization, regardless of application rate.

Cumulative N_2_O emissions increased immediately at the time of fertilizer application and were 80% nitrification-derived by mid-June 2019 (Figure S5). However, for a given fertilizer application rate, cumulative annual N_2_O emissions converged across all three timing experiments by the end of the growing season. Increases in denitrification-derived N_2_O emissions compensated for reductions in nitrification-derived N_2_O emissions earlier in the growing season (Figure S5). Conditions suitable for denitrification increased in the latter part of the 2019 growing season as soil moisture increased whereas soil moisture was lowest during July and August (Figure S6). By the end of 2019, 78% of simulated N_2_O emissions were nitrification-derived and 22% were denitrification-derived.

### Sorghum legacy effects on soybean nitrogen losses

Although no nitrification suppression or fertilizer application was simulated during the soybean years (2019 and 2020), we observed residual effects from the sorghum years (2018 and 2019) on N losses from soybean (Figure 1; Table S6). N_2_O emissions, NO_3_^-^ leaching, and gross nitrification all increased with the previous year’s fertilization rate (Figure 1). However, there were only small effects of nitrification suppression from the previous year on either N_2_O emissions, NO_3_^-^ leaching, or gross nitrification (Figure 1). Across all 2019 sorghum fertilization rates, 2020 soybean N_2_O emissions were stimulated by up to 0.2% in the measNS experiment and up to 2% in the maxNS experiment (Table S6). In contrast, 2020 soybean NO_3_^-^ leaching was consistently reduced in the measNS and maxNS experiments by 0.2-0.6% and 0.6-3.0%, respectively (Table S6).

Altering the timing of sorghum fertilizer application in 2019 generally reduced N_2_O emissions and NO_3_^-^ leaching in 2020 soybean. Soil N_2_O emissions were reduced by 0.5 to 3% relative to fertilizer addition at planting in all but the 168 kg N ha^-1^ y^-1^ split fertilizer addition. Nitrate leaching was reduced by 2-6% across all fertilizer application rates in all three fertilizer timing experiments.

### Long-term effects of nitrification suppression and fertilizer timing on nitrogen losses

Over the period of 2019-2030, with sorghum simulated in odd years and soybean simulated with no fertilization or nitrification suppression in even years, nitrification suppression reduced cumulative N losses by a small amount. The measNS experiment showed a 0.5-1% reduction in cumulative N_2_O emissions and <1% reductions in cumulative NO_3_^-^ leaching relative to the noNS experiment (Table 2). In contrast, the maxNS experiment showed a 1.5-5% reduction in cumulative N_2_O emissions and a 1-3% reduction in cumulative NO_3_^-^ leaching compared to the same fertilizer application rate in the None experiment; the N loss reductions increased with application rate (Table 2).

Fertilizer timing experiments showed mixed effects on cumulative 2019–2030 N losses. The 3-week and 5-week delayed experiments showed changes from -2% to +2% in cumulative N_2_O emissions relative to no fertilizer delay at the same fertilizer application rate (Table 2). For fertilizer application rates of 56 and 112 kg N ha^-1^ y^-1^, the delayed experiments reduced total nitrate leaching 2–4% relative to no fertilizer delay; however, total nitrate leaching increased by 16–18% when fertilizer application rate was 168 kg N ha^-1^ y^-1^ (Table 2). The split fertilizer experiment showed a 1-4% reduction in cumulative N_2_O emissions and a 2-4% reduction in cumulative NO_3_^-^ leaching compared to the same fertilizer application rate in the no delay experiment (Table 2).

Changes in 2019-2030 cumulative N_2_O and NO_3_^-^ leaching from altering fertilizer N application rates were a magnitude greater than in any other N loss mitigation experiment (Table 2). Compared to corresponding experiments with no UAN fertilizer, experiments with 56 kg N ha^-1^ y^-1^ fertilization increased N_2_O emissions by 18-22% and cumulative NO_3_^-^ leaching by 13–17%; experiments with 112 kg N ha^-1^ y^-1^ fertilization increased N_2_O emissions by 45-50% and cumulative NO_3_^-^ leaching by 31-37%; experiments with 168 kg N ha^-1^ y^-1^ fertilization increased N_2_O emissions by 81–93% and cumulative NO_3_^-^ leaching by 65–100%.

## Discussion

We hypothesized that the effectiveness of nitrification suppression in reducing agroecosystem N losses via soil N_2_O emissions and NO_3_^-^ leaching would depend on the synchronization between nitrification suppression and the fertilizer-induced spike in nitrification. As expected, our model experiments showed that, with conventional timing of fertilizer application with planting, nitrification suppression led to relatively low (< 9%) reductions in annual N losses for energy sorghum. Consistent with rapid nitrification of NH_4_^+^ applied in fertilizer (Norton and Ouyang 2019), gross nitrification rates peaked before nitrification suppression started in the fifth week after planting (Figure 3). Our fertilizer timing model experiments showed that, as expected, delayed and split fertilizer application delayed the fertilizer-induced spike in nitrification to better synchronize it with measured seasonal patterns in nitrification suppression levels. However, we found that the effect of fertilizer timing on gross nitrification and N loss rates depended on temperature and soil moisture around the time of fertilizer application such that these rates could increase or decrease, depending on the weather in a given year. Moreover, we found greater effects of nitrification suppression on annual rates of soil N_2_O emissions than NO_3_^-^ leaching, likely due to differing environmental controls on these two N loss pathways. Here we discuss potential mechanisms leading to these patterns in nitrification suppression and fertilizer timing effects predicted by the DayCent-CABBI model and the implications of our findings for the potential use of nitrification suppression to reduce agroecosystem N losses in the Midwest U.S. and elsewhere.

Nitrification suppression may have little effect on annual cumulative gross nitrification rates regardless of the timing of nitrification suppression or fertilizer application. Our model experiments show that delaying or splitting fertilizer application reduces early growing season gross nitrification rates by shifting later the fertilizer-induced spike in nitrification and by synchronizing the spike with start of nitrification suppression to reduce the magnitude of the spike (Figure 4). However, this reduction is offset by increased gross nitrification rates later in the growing season, likely by shifting elevated soil NH_4_^+^ concentrations to when warmer summer temperatures stimulate microbial activity in general (Agehara and Warncke 2005, Ouyang et al. 2017, Norton and Ouyang 2019). Therefore, altering fertilizer timing had little effect on cumulative gross nitrification rates at the annual time scale (Figure 2EF). In irrigated systems, higher moisture and temperature later in the season could further increase annual gross nitrification when fertilization is delayed (Agehara and Warncke 2005). Similar to our findings, altering fertilizer timing had inconsistent effects on losses of both NO_3_^-^ leaching (Eagle et al. 2017) and N_2_O emissions (Zebarth et al. 2008, Phillips et al. 2009, Zebarth et al. 2012, Han et al. 2017) in past field studies. These results suggest that environmental controls on nitrification should be taken into account when considering potential effects of nitrification suppression and fertilizer timing on annual cumulative gross nitrification rates.

Simulated effects of nitrification suppression and fertilizer timing on N_2_O emissions mirrored effects on gross nitrification. This is largely due to the predicted role of nitrification as the dominant N_2_O source at our site and the source of NO_3_^-^ for denitrification. Our model experiments show that nitrification is responsible for 78% of annual soil N_2_O emissions, consistent with some empirical studies in cropping systems (Skiba et al. 1993, Bremner 1997, Stevens et al. 1997, Toyoda et al. 2011, Tierling and Kuhlmann 2018) but inconsistent with other empirical studies showing denitrification as the dominant N_2_O source (Müller et al. 2014, Li et al. 2016, Liang and Robertson 2021). In particular, nitrification appears to be a significant source of N_2_O when NH_4_^+^ substrate is abundant and soils are not fully saturated (Bremner 1997, Toyoda et al. 2011), which is consistent with the significant increase in both gross nitrification and N_2_O emissions after fertilization in our study. If N_2_O is mostly nitrification-derived after fertilization, delaying fertilizer addition to better align N availability with the suppression of nitrification should lead to reduced N_2_O losses (Ferrari Machado et al. 2020). However, because delayed or split fertilizer timing also led to high soil NO_3_^-^ concentrations later in the growing season when soil moisture conditions were more conducive for denitrification (Bremner 1997) (Figure S6), greater late season denitrification-derived soil N_2_O emissions partially offset earlier season reductions in nitrification-derived soil N_2_O emissions (Figure S5). Therefore, reductions in annual soil N_2_O emissions due to nitrification suppression or changes in fertilizer timing were smaller than reductions in cumulative annual gross nitrification rates.

Our model experiments suggest that NO_3_^-^ leaching is dominantly controlled by water flux such that changes in soil NO_3_^-^ concentrations caused by nitrification suppression or different fertilizer management have limited effects on annual NO_3_^-^ leaching rates. In the Midwestern US, a large proportion of annual NO_3_^-^ leaching occurs in the non-growing season, especially April-June, when low evapotranspiration rates lead to substantial water flux through the soil profile and high rates of NO_3_-loss out of the soil (Di and Cameron 2002a, Randall and Vetsch 2005, Liao et al. 2012, Hess et al. 2020). Therefore, suppression of nitrification, either by plants or synthetic nitrification inhibitors applied in-season (Eagle et al. 2017), has a limited impact on annual NO_3_^-^ leaching rates. However, in regions with high precipitation mid-growing season or in irrigated systems, NO_3_^-^ losses could be temporally synchronized with nitrification suppression (Ritter 1989, Nyamangara et al. 2003, Mack et al. 2005, van der Laan et al. 2010), and since nitrification increases with temperature and moisture (Agehara and Warncke 2005), suppression by plants could have a larger effect. Surprisingly, the model predicted little to no effect of fertilizer application rate on NO_3_^-^ leaching in 2018 or 2019 (Figure 2EF), which is inconsistent with past studies in the Midwest showing increased N leaching with higher fertilizer application rates (Helmers et al. 2012, Smith et al. 2013, Hussain et al. 2020). In our study, it appears that the rate of fertilizer addition had a larger impact on NO_3_^-^ leaching the following growing season. For example, despite a lack of fertilizer effect in 2018 sorghum plots, higher fertilization in 2018 increased the amount of NO_3_^-^ leached the following season then the plots were planted with soybean (Figure 1). We also found a net positive effect of fertilization rate on NO_3_^-^ leaching from 2019-2030, which accounts for lag effects in the long-term (Table 2). In addition, the recommended fertilization rate for bioenergy sorghum is 112 kg N ha^-1^, and we detected significant increases in NO_3_^-^ leaching when we exceeded this recommended rate and applied 168 kg N ha^-1^ (Table 2). However, there are also challenges in measuring the field-scale effects of management practices on NO_3_^-^ exiting the soil profile (Hess et al. 2020, Wey et al. 2021), so empirical datasets for calibrating and validating NO_3_^-^ leaching in models are scarce. Since there is significant water flux through the soil profile during the non-growing season after sorghum harvest and prior to crop establishment the next year, our results underscore the importance of water flux and variability in precipitation in controlling NO_3_^-^ leaching, especially during fallow periods (Di and Cameron 2002a, Randall and Vetsch 2005, Liao et al. 2012).

Since nitrification suppression by sorghum had only very small effects on N losses, we also found no evidence of legacy effects of sorghum nitrification suppression on N losses in the following soybean growing seasons. In fact, despite low leaching rates of NO_3_^-^ during the 2018 sorghum growing season, much of the NO_3_^-^ that had been stored within the soil was lost during the wetter 2019 soybean growing season (Figure 1). When sorghum-soybean rotations were simulated through 2030 over a realistic range of weather conditions, consistent but small reductions in long-term cumulative N losses (Table 2) illustrate that sorghum nitrification suppression does have some control over N_2_O emissions and leaching of NO_3_^-^. However, the limited magnitude of sorghum’s influence further emphasizes the strength of environmental control and impact of mineralization on N losses (Di and Cameron 2002a, Loecke et al. 2017, Leitner et al. 2020). In addition, reducing the amount of N fertilizer applied was more effective in reducing N_2_O emissions and nitrate leaching over the long-term than in 2018 and 2019 model simulations, and reducing N inputs had a much stronger effect than any rate of nitrification suppression.

Management strategies targeted at retaining N in the non-growing season may be more effective for reducing annual N losses from agroecosystems in climates such as the rainfed Midwest where spring thaw and fall precipitation drive a large proportion of annual NO_3_^-^ leaching. For example, incorporating winter cover crops into annual rotations eliminates the spring fallow period and is an effective strategy for limiting spring N losses (Di and Cameron 2002a, Jewett and Thelen 2007, Blanco-Canqui et al. 2021). Since bioenergy and biomass sorghum crops are harvested for aboveground biomass rather than grain yield, farmers have more flexibility in the timing of harvest and cover crop planting in the late summer or early fall when sorghum biomass begins to plateau (Schetter et al. 2021). This could provide a unique opportunity for synergy between nitrification suppression by sorghum during the growing season and uptake of available N during the fall and spring by a cover crop. Combining nitrification suppression during the growing season with off-season management to retain N is likely to achieve lower N losses and better agroecosystem sustainability than either strategy alone.

## Conclusion

Despite recent advances in our understanding of the ability of some plant species to suppress nitrification in the rhizosphere, the impact on agroecosystem N losses has remained poorly characterized. We show that nitrification suppression in the rhizosphere of energy sorghum has only a limited capacity to reduce field-scale emissions of N_2_O and leaching of NO_3_^-^ due to asynchrony between peak gross nitrification in the spring prior to crop establishment and suppression of nitrification by sorghum plants mid-season. Thus, breeding or engineering crops to simply increase nitrification suppression will lead to only modest reductions in N losses in rainfed regions where significant precipitation occurs outside of the growing season. Altering the timing of fertilizer addition to better synchronize N availability with crop N demand and nitrification suppression can reduce N losses. However, delaying fertilization also increases N availability when environmental conditions, especially soil temperature and moisture, favor higher biological activity (i.e., nitrification and denitrification rates), reducing and sometimes negating the effect of altered fertilizer timing on cumulative N losses. Additionally, because the controls on N_2_O emissions and NO_3_^-^ leaching are different, the magnitude and direction of effects of nitrification suppression and fertilizer timing on these two N loss pathways can differ. Overall, management strategies targeted at retaining N in the non-growing season may be more effective for reducing annual N losses in the rainfed Midwest, particularly for NO_3_^-^ leaching in the early spring. Given the potential for different timing of N losses in irrigated systems, we suggest that the effects of nitrification suppression on N_2_O emissions and NO_3_^-^ leaching be further explored in regions where irrigation is more common. Further research on nitrification suppression should prioritize optimizing its timing and magnitude alongside off-season N retention strategies to improve the N sustainability of agroecosystems.

## Acknowledgements

The authors thank Adam von Haden and Ryan Quinn for their efforts quantifying root biomass in these trials. We also thank August Schetter and DK Lee for measuring and providing aboveground biomass and yield data. We also thank Avarna Jain, Taylor Bozman, Allison Cook, Ingrid Holstrom, Rachel van Allen, Elle Lucadamo, Sierra Raglin, Rachel Waltermire, and Sandra Simon for their help with measurements of soil N pools and nitrification suppression. We also thank Dr. Bill Rooney for supplying the germplasm via the Texas A&M Sorghum Breeding Program. Finally, we thank Tim Mies and Trace Elliott for their management of planting and fertilization at the UIUC Energy Farm. This work was supported by the DOE Center for Advanced Bioenergy and Bioproducts Innovation (U.S. Department of Energy, Office of Science, Office of Biological and Environmental Research under Award Number DE-SC0018420).

## Author contributions

MDH led the writing of the manuscript and performed all DayCent model simulations. MB and AF provided field data. All authors contributed critically to the draft and gave final approval for publication.

## Notes

### Competing Interest Statement

The authors have declared no competing interest.

## References

Agehara, S., and D. D. Warncke. 2005. Soil Moisture and Temperature Effects on Nitrogen Release from Organic Nitrogen Sources. Soil Science Society of America Journal 69:1844–1855.

Belser, L. W., and E. L. Mays. 1980. Sepcific inhibition of nitrite oxidation by chlorate and its use in assessing nitrification in soils and sediments. Applied and Environmental Microbiolosy 39:505–510.

Benjamin, J. G., and D. C. Nielsen. 2004. A method to separate plant roots from soil and analyze root surface area. Plant and Soil 267:225–234.

Black, C. K., M. D. Masters, D. S. LeBauer, K. J. Anderson-Teixeira, and E. H. DeLucia. 2017. Root volume distribution of maturing perennial grasses revealed by correcting for minirhizotron surface effects. Plant and Soil 419:391–404.

Blanco-Canqui, H., S. J. Ruis, J. D. Holman, C. F. Creech, and A. K. Obour. 2021. Can cover crops improve soil ecosystem services in water-limited environments? A review. Soil Science Society of America Journal 86:1–18.

Booth, M. S., J. M. Stark, and E. Rastetter. 2005. Controls on Nitrogen Cycling in Terrestrial Ecosystems: A Synthetic Analysis of Literature Data. Ecological Monographs 75:139–157.

Bremner, J. M. 1997. Sources of nitrous oxide in soils. Nutrient Cycling in Agroecosystems 49:7–16.

Burnham, M. B., S. J. Simon, D. Lee, A. D. Kent, E. H. DeLucia, and W. H. Yang. 2022. Intra-and inter-annual variability of nitrification in the rhizosphere of field-grown bioenergy sorghum. GCB Bioenergy 14:393–410.

Cao, P., C. Lu, and Z. Yu. 2018. Historical nitrogen fertilizer use in agricultural ecosystems of the contiguous United States during 1850–2015: application rate, timing, and fertilizer types. Earth System Science Data 10:969–984.

Chaparro, J. M., D. V. Badri, M. G. Bakker, A. Sugiyama, D. K. Manter, and J. M. Vivanco. 2013. Root exudation of phytochemicals in Arabidopsis follows specific patterns that are developmentally programmed and correlate with soil microbial functions. PLoS One 8.

Ciampitti, I. A., and T. J. Vyn. 2011. A comprehensive study of plant density consequences on nitrogen uptake dynamics of maize plants from vegetative to reproductive stages. Field Crops Research 121:2–18.

Coskun, D., D. T. Britto, W. Shi, and H. J. Kronzucker. 2017. Nitrogen transformations in modern agriculture and the role of biological nitrification inhibition. Nat Plants 3:17074.

Dai, S., M. D. Shulski, K. G. Hubbard, and E. S. Takle. 2016. A spatiotemporal analysis of Midwest US temperature and precipitation trends during the growing season from 1980 to 2013. International Journal of Climatology 36:517–525.

Del Grosso, S., A. Mosier, W. Parton, and D. Ojima. 2005. DAYCENT model analysis of past and contemporary soil NO and net greenhouse gas flux for major crops in the USA. Soil and Tillage Research 83:9–24.

Del Grosso, S., D. Ojima, W. Parton, A. Mosier, G. Peterson, and D. Schimel. 2002. Simulated effects of dryland cropping intensification on soil organic matter and greenhouse gas exchanges using the DAYCENT ecosystem model. Environmental Pollution 116:S75–S83.

Del Grosso, S. J., S. M. Ogle, W. J. Parton, and F. J. Breidt. 2010. Estimating uncertainty in N2O emissions from U.S. cropland soils. Global Biogeochemical Cycles 24:/a-n/a.

Di, H. J., and K. C. Cameron. 2002a. Nitrate leaching in temperate agroecosystems: sources, factors and mitigating strategies. Nutrient Cycling in Agroecosystems 64:237–256.

Di, H. J., and K. C. Cameron. 2002b. The use of a nitrification inhibitor, dicyandiamide (DCD), to decrease nitrate leaching and nitrous oxide emissions in a simulated grazed and irrigated grassland. Soil Use and Management 18:395–403.

Duval, B. D., M. Hartman, E. Marx, W. J. Parton, S. P. Long, and E. H. DeLucia. 2015. Biogeochemical consequences of regional land use change to a biofuel crop in the southeastern United States. Ecosphere 6:art265.

Eagle, A. J., L. P. Olander, K. L. Locklier, J. B. Heffernan, and E. S. Bernhardt. 2017. Fertilizer management and environmental factors drive N2O and NO3 losses in corn: A meta-analysis. Soil Science Society of America Journal 81:1191–1202.

Elrys, A. S., J. Wang, M. A. S. Metwally, Y. Cheng, J. B. Zhang, Z. C. Cai, S. X. Chang, and C. Muller. 2021. Global gross nitrification rates are dominantly driven by soil carbon-to-nitrogen stoichiometry and total nitrogen. Glob Chang Biol 27:6512–6524.

Epa, U. 2018. Greenhouse gas emissions and sinks, 1990-2016. in U. Epa, editor.

Farquharson, R. 2016. Nitrification rates and associated nitrous oxide emissions from agricultural soils – a synopsis. Soil Research 54:469.

Ferrari Machado, P. V., K. Neufeld, S. E. Brown, P. R. Voroney, T. W. Bruulsema, and C. Wagner-Riddle. 2020. High temporal resolution nitrous oxide fluxes from corn (Zea mays L.) in response to the combined use of nitrification and urease inhibitors. Agriculture, Ecosystems & Environment 300:106996.

Finzi, A. C., R. Z. Abramoff, K. S. Spiller, E. R. Brzostek, B. A. Darby, M. A. Kramer, and R. P. Phillips. 2015. Rhizosphere processes are quantitatively important components of terrestrial carbon and nutrient cycles. Glob Chang Biol 21:2082–2094.

Gilsanz, C., D. Báez, T. H. Misselbrook, M. S. Dhanoa, and L. M. Cárdenas. 2016. Development of emission factors and efficiency of two nitrification inhibitors, DCD and DMPP. Agriculture, Ecosystems & Environment 216:1–8.

Han, Z., M. T. Walter, and L. E. Drinkwater. 2017. N2O emissions from grain cropping systems: a meta-analysis of the impacts of fertilizer-based and ecologically-based nutrient management strategies. Nutrient Cycling in Agroecosystems 107:335–355.

Helmers, M. J., X. Zhou, J. L. Baker, S. W. Melvin, and D. W. Lemke. 2012. Nitrogen loss on tile-drained Mollisols as affected by nitrogen application rate under continuous corn and corn-soybean rotation systems. Canadian Journal of Soil Science 92:493–499.

Hess, L. J. T., E.-L. S. Hinckley, G. P. Robertson, and P. A. Matson. 2020. Rainfall intensification increases nitrate leaching from tilled but not no-till cropping systems in the U.S. Midwest. Agriculture, Ecosystems & Environment 290:106747.

Huff, F. A., and J. R. Angel. 1992. Rainfall atlas of the Midwest. Illinois State Water Survey, Champaign, IL.

Hussain, M. Z., G. P. Robertson, B. Basso, and S. K. Hamilton. 2020. Leaching losses of dissolved organic carbon and nitrogen from agricultural soils in the upper US Midwest. Sci Total Environ 734:139379.

Inselsbacher, E., N. Hinko-Najera Umana, F. C. Stange, M. Gorfer, E. Schüller, K. Ripka, S. Zechmeister-Boltenstern, R. Hood-Novotny, J. Strauss, and W. Wanek. 2010. Short-term competition between crop plants and soil microbes for inorganic N fertilizer. Soil Biology and Biochemistry 42:360–372.

Janke, C. K., L. A. Wendling, and R. Fujinuma. 2018. Biological nitrification inhibition by root exudates of native species, Hibiscus splendens and Solanum echinatum. PeerJ 6:e4960.

Jewett, M. R., and K. D. Thelen. 2007. Winter Cereal Cover Crop Removal Strategy Affects Spring Soil Nitrate Levels. Journal of Sustainable Agriculture 29:55–67.

Kent, J., M. D. Hartman, D. K. Lee, and T. Hudiburg. 2020. Simulated Biomass Sorghum GHG Reduction Potential is Similar to Maize. Environ Sci Technol.

Leitner, S., T. Dirnbock, J. Kobler, and S. Zechmeister-Boltenstern. 2020. Legacy effects of drought on nitrate leaching in a temperate mixed forest on karst. J Environ Manage 262:110338.

Li, X., P. Sørensen, J. E. Olesen, and S. O. Petersen. 2016. Evidence for denitrification as main source of N 2 O emission from residue-amended soil. Soil Biology and Biochemistry 92:153–160.

Liang, D., and G. P. Robertson. 2021. Nitrification is a minor source of nitrous oxide (N2 O) in an agricultural landscape and declines with increasing management intensity. Glob Chang Biol 27:5599–5613.

Liao, L., C. T. Green, B. A. Bekins, and J. K. Böhlke. 2012. Factors controlling nitrate fluxes in groundwater in agricultural areas. Water Resources Research 48.

Loecke, T. D., A. J. Burgin, D. A. Riveros-Iregui, A. S. Ward, S. A. Thomas, C. A. Davis, and M. A. S. Clair. 2017. Weather whiplash in agricultural regions drives deterioration of water quality. Biogeochemistry 133:7–15.

Mack, U. D., K. H. Feger, Y. Gong, and K. Stahr. 2005. Soil water balance and nitrate leaching in winter wheat–summer maize double-cropping systems with different irrigation and N fertilization in the North China Plain. Journal of Plant Nutrition and Soil Science 168:454–460.

Moll, R. H., E. J. Kamprath, and W. A. Jackson. 1982. Analysis and Interpretation of Factors Which Contribute to Efficiency of Nitrogen Utilization. Agronomy Journal 74:562–564.

Moore, C. E., D. M. Berardi, E. Blanc-Betes, E. C. Dracup, S. Egenriether, N. Gomez-Casanovas, M. D. Hartman, T. Hudiburg, I. Kantola, M. D. Masters, W. J. Parton, R. Van Allen, A. C. Haden, W. H. Yang, E. H. DeLucia, and C. J. Bernacchi. 2020a. The carbon and nitrogen cycle impacts of reverting perennial bioenergy switchgrass to an annual maize crop rotation. GCB Bioenergy 12:941–954.

Moore, C. E., A. C. Haden, M. B. Burnham, I. B. Kantola, C. D. Gibson, B. J. Blakely, E. C. Dracup, M. D. Masters, W. H. Yang, E. H. DeLucia, and C. J. Bernacchi. 2020b. Ecosystem-scale biogeochemical fluxes from three bioenergy crop candidates: How energy sorghum compares to maize and miscanthus. GCB Bioenergy 13:445–458.

Müller, C., R. J. Laughlin, O. Spott, and T. Rütting. 2014. Quantification of N2O emission pathways via a 15N tracing model. Soil Biology and Biochemistry 72:44–54.

Nardi, P., H. J. Laanbroek, G. W. Nicol, G. Renella, M. Cardinale, G. Pietramellara, W. Weckwerth, A. Trinchera, A. Ghatak, and P. Nannipieri. 2020. Biological nitrification inhibition in the rhizosphere: determining interactions and impact on microbially mediated processes and potential applications. FEMS Microbiol Rev 44:874–908.

Norton, J., and Y. Ouyang. 2019. Controls and Adaptive Management of Nitrification in Agricultural Soils. Front Microbiol 10:1931.

Nyamangara, J., L. F. Bergstrom, M. I. Piha, and K. E. Giller. 2003. Fertilizer use efficiency and nitrate leaching in a tropical sandy soil. J Environ Qual 32:599–606.

O’Sullivan, C. A., I. R. P. Fillery, M. M. Roper, and R. A. Richards. 2016. Identification of several wheat landraces with biological nitrification inhibition capacity. Plant and Soil 404:61–74.

Ogle, S. M., F. J. Breidt, M. Easter, S. Williams, K. Killian, and K. Paustian. 2010. Scale and uncertainty in modeled soil organic carbon stock changes for US croplands using a process-based model. Global Change Biology 16:810–822.

Ouyang, Y., J. M. Norton, and J. M. Stark. 2017. Ammonium availability and temperature control contributions of ammonia oxidizing bacteria and archaea to nitrification in an agricultural soil. Soil Biology and Biochemistry 113:161–172.

Parton, W. J., M. Hartman, D. Ojima, and D. Schimel. 1998. DAYCENT and its land surface submodel: description and testing. Global and Planetary Change 19:35–48.

Phillips, R. L., D. L. Tanaka, D. W. Archer, and J. D. Hanson. 2009. Fertilizer application timing influences greenhouse gas fluxes over a growing season. J Environ Qual 38:1569–1579.

Quinn, R. 2021. Quantifying and comparing belowground carbon pools and fluxes of two bioenergy crop species: Miscanthus x giganeus and Sorghum bicolor. Boston University.

Randall, G. W., and J. A. Vetsch. 2005. Nitrate losses in subsurface drainage from a corn-soybean rotation as affected by fall and spring application of nitrogen and nitrapyrin. J Environ Qual 34:590–597.

Ritter, W. F. 1989. Nitrate leaching under irrigation in the United States—a review. Journal of Environmental Science and Health. Part A: Environmental Science and Engineering 24:349–378.

Sarr, P. S., Y. Ando, S. Nakamura, S. Deshpande, and G. V. Subbarao. 2019. Sorgoleone release from sorghum roots shapes the composition of nitrifying populations, total bacteria, and archaea and determines the level of nitrification. Biology and Fertility of Soils 56:145–166.

Schetter, A., C.-H. Lin, C. Zumpf, C. Jang, L. Hoffmann, W. Rooney, and D. K. Lee. 2021. Genotype-Environment-Management Interactions in Biomass Yield and Feedstock Composition of Photoperiod-Sensitive Energy Sorghum. BioEnergy Research.

Six, J., S. M. Ogle, F. Jay breidt, R. T. Conant, A. R. Mosier, and K. Paustian. 2004. The potential to mitigate global warming with no-tillage management is only realized when practised in the long term. Global Change Biology 10:155–160.

Skiba, U., K. A. Smith, and D. fowler. 1993. Nitrification and denitrification as sources of nitric oxide and nitrous oxide in a sandy loam soil. Soil Biology and Biochemistry 25:1527–1536.

Smith, C. M., M. B. David, C. A. Mitchell, M. D. Masters, K. J. Anderson-Teixeira, C. J. Bernacchi, and E. H. Delucia. 2013. Reduced nitrogen losses after conversion of row crop agriculture to perennial biofuel crops. J Environ Qual 42:219–228.

Stevens, R. J., R. J. Laughlin, L. C. Burns, J. R. M. Arah, and R. C. Hood. 1997. Measuring the contributions of nitrification and denitrification to the flux of nitrous oxide from soil. Soil Biology and Biochemistry 29:139–151.

Stienstra, A. W., P. Klein Gunnewiek, and H. J. Laanbroek. 1994. Repression of nitrification in soils under a climax grassland vegetation. FEMS Microbiology Ecology 14:45–52.

Straube, J. R., M. Chen, W. J. Parton, S. Asso, Y.-A. Liu, D. S. Ojima, and W. Gao. 2018. Development of the DayCent-Photo model and integration of variable photosynthetic capacity. Frontiers of Earth Science 12:765–778.

Subbarao, G. V., M. Kishii, A. Bozal-Leorri, I. Ortiz-Monasterio, X. Gao, M. I. Ibba, H. Karwat, M. B. Gonzalez-Moro, C. Gonzalez-Murua, T. Yoshihashi, S. Tobita, V. Kommerell, H. J. Braun, and M. Iwanaga. 2021. Enlisting wild grass genes to combat nitrification in wheat farming: A nature-based solution. Proc Natl Acad Sci U S A 118.

Subbarao, G. V., K. Nakahara, T. Ishikawa, H. Ono, M. Yoshida, T. Yoshihashi, Y. Zhu, H. A. K. M. Zakir, S. P. Deshpande, C. T. Hash, and K. L. Sahrawat. 2013a. Biological nitrification inhibition (BNI) activity in sorghum and its characterization. Plant and Soil 366:243–259.

Subbarao, G. V., K. L. Sahrawat, K. Nakahara, I. M. Rao, M. Ishitani, C. T. Hash, M. Kishii, D. G. Bonnett, W. L. Berry, and J. C. Lata. 2013b. A paradigm shift towards low-nitrifying production systems: the role of biological nitrification inhibition (BNI). Ann Bot 112:297–316.

Teutscherová, N., E. Vázquez, J. Trubač, D. M. Villegas, G. V. Subbarao, M. Pulleman, and J. Arango. 2021. Gross N transformation rates in soil system with contrasting Urochloa genotypes do not confirm the relevance of BNI as previously assessed in vitro. Biology and Fertility of Soils.

Tierling, J., and H. Kuhlmann. 2018. Emissions of nitrous oxide (N2O) affected by pH-related nitrite accumulation during nitrification of N fertilizers. Geoderma 310:12–21.

Toyoda, S., M. Yano, S.-i. Nishimura, H. Akiyama, A. Hayakawa, K. Koba, S. Sudo, K. Yagi, A. Makabe, Y. Tobari, N. O. Ogawa, N. Ohkouchi, K. Yamada, and N. Yoshida. 2011. Characterization and production and consumption processes of N2O emitted from temperate agricultural soils determined via isotopomer ratio analysis. Global Biogeochemical Cycles 25:n/a-n/a.

van der Laan, M., R. J. Stirzaker, J. G. Annandale, K. L. Bristow, and C. C. d. Preez. 2010. Monitoring and modelling draining and resident soil water nitrate concentrations to estimate leaching losses. Agricultural Water Management 97:1779–1786.

Van Meter, K. J., N. B. Basu, and P. Van Cappellen. 2017. Two centuries of nitrogen dynamics: Legacy sources and sinks in the Mississippi and Susquehanna River Basins. Global Biogeochemical Cycles 31:2–23.

van Oosterom, E. J., A. K. Borrell, S. C. Chapman, I. J. Broad, and G. L. Hammer. 2010. Functional dynamics of the nitrogen balance of sorghum: I. N demand of vegetative plant parts. Field Crops Research 115:19–28.

Wey, H., D. Hunkeler, W. A. Bischoff, and E. K. Bunemann. 2021. Field-scale monitoring of nitrate leaching in agriculture: assessment of three methods. Environ Monit Assess 194:4.

Yang, M., Y. Fang, D. Sun, and Y. Shi. 2016. Efficiency of two nitrification inhibitors (dicyandiamide and 3, 4-dimethypyrazole phosphate) on soil nitrogen transformations and plant productivity: a meta-analysis. Sci Rep 6:22075.

Zakir, H. A., G. V. Subbarao, S. J. Pearse, S. Gopalakrishnan, O. Ito, T. Ishikawa, N. Kawano, K. Nakahara, T. Yoshihashi, H. Ono, and M. Yoshida. 2008. Detection, isolation and characterization of a root-exuded compound, methyl 3-(4-hydroxyphenyl) propionate, responsible for biological nitrification inhibition by sorghum (Sorghum bicolor). New Phytol 180:442–451.

Zebarth, B. J., P. Rochette, D. L. Burton, and M. Price. 2008. Effect of fertilizer nitrogen management on N2O emissions in commercial corn fields. Canadian Journal of Soil Science 88:189–195.

Zebarth, B. J., E. Snowdon, D. L. Burton, C. Goyer, and R. Dowbenko. 2012. Controlled release fertilizer product effects on potato crop response and nitrous oxide emissions under rain-fed production on a medium-textured soil. Canadian Journal of Soil Science 92:759–769.

