## Supplemental Materials for "In Silico Evaluation of Plant Nitrification Suppression Effects on Agroecosystem Nitrogen Loss"

Table S1. Agricultural history (2010-2020) for the 2018 and 2019 sorghum plots.

| Agricultural history of sorghum 2018 plots | Agricultural history of sorghum 2019 plots |
| --- | --- |
| 2010: Soybean | 2010: Soybean |
| 2011: Miscanthus Trials | 2011: Sorghum (UAN 101 kg N $ha^{-1}$) |
| 2012: Miscanthus Trials | 2012: Soybean |
| 2013: Miscanthus Trials | 2013: Sorghum (UAN 101 kg N $ha^{-1}$) |
| 2014: Miscanthus Trials | 2014: Soybean |
| 2015: Miscanthus Trials | 2015: Sorghum (UAN 101 kg N $ha^{-1}$) |
| 2016: Corn (UAN 202 kg N $ha^{-1}$) | 2016: Soybean |
| 2017: Soybean | 2017: Corn (UAN 202 kg N $ha^{-1}$) |
| **2018: Sorghum NS (UAN 0,56,112,168 kg N $ha^{-1}$)** | 2018: Soybean |
| 2019: Soybean | **2019: Sorghum NS (UAN 0,56,112,168 kg N $ha^{-1}$)** |
| | 2020: Soybean |

Table S2. Simplified DayCent schedule file block for sorghum 2018 and sorghum 2019 plots. Nitrification suppressions (NS) is simulated as a fertilizer event where no N is applied but nitrification inhibitors are applied with the specified rate of reduction of gross nitrification. NS was measured three times each sorghum year, and for simulation purposes NS rates were interpolated between the measured rates.

| **Sorghum 2018 followed by soybean 2019** | | **Sorghum 2019 preceded by soybean 2018** | |
|---|---|---|---|
| **Year day event** | **# Comment** | **Year day event** | **# Comment** |
| 2018 138 CROP SORG | # plant sorghum May-18 | 2018 139 CROP SYBN | # plant soybean May-19 |
| 2018 142 FERT UAN | # fertilize May-22 | 2018 285 HARV | # harvest soybean |
| 2018 178 FERT NS03 | # NS 3% Jun-27 | 2018 285 LAST | # end-of-growing season |
| 2018 192 FERT NS12 | | 2019 152 CROP SORG | # plant sorghum Jun-1 |
| 2018 206 FERT NS20 | # NS 20% Jul-25 | 2019 158 FERT UAN | # fertilize Jun-7 |
| 2018 220 FERT NS12 | | 2019 189 FERT NS14 | # NS 14% Jul-8 |
| 2018 234 FERT NS05 | # NS 5% Aug-22 | 2019 203 FERT NS36 | |
| 2018 288 HARV | # harvest sorghum Oct-15 | 2019 218 FERT NS58 | # NS 58% Aug-5 |
| 2018 288 LAST | # end-of-growing season | 2019 232 FERT NS40 | |
| 2019 139 CROP SYBN | # plant soybean May-19 | 2019 246 FERT NS22 | # NS 22% Sep-3 |
| 2019 285 HARV | # harvest soybean | 2019 283 HARV | # harvest sorghum Oct-9 |
| 2019 285 LAST | # end-of-growing season | 2019 283 LAST | # end-of-growing season |

Table S3. Measured ammonium and nitrate levels, pre- and post-growing season compared to simulated values.

| **Date** | **measured $NH_4^+$ (kg N $ha^{-1}$)** | **simulated $NH_4^+$ (kg N $ha^{-1}$)** | **measured $NO_3^-$ (kg N $ha^{-1}$)** | **simulated $NO_3^-$ (kg N $ha^{-1}$)** |
|---|---|---|---|---|
| May 18, 2018 (pre-fertilization) | 0.8 (± 0.3) (0-15cm) | 2.9 (total $NH_4^+$ pool) | 14.6 (± 1.3) (0-15cm) | 7.4 (0-15cm)<br>18.0 (0-30cm) |
| June 7, 2019 (pre-fertilization) | 2.2 (± 0.2) (0-30cm) | 3.6 (total $NH_4^+$ pool) | 25.6 (± 0.7) (0-30cm) | 23.0 (0-30 cm) |
| October 1, 2018 | 25.0 (± 1.7) (0-50 cm) | 2.0 (total $NH_4^+$ pool) | 25.0 (± 7.6) (0-50 cm) | 1.2 (0-45 cm)<br>14.3 (0-105 cm) |

Table S4. Simulated $N_2O$ emissions and nitrate leaching for sorghum 2019 comparing nitrification suppression experiments (noNS = no nitrification suppression, measNS = measured nitrification suppression, maxNS = maximum nitrification suppression) and fertilizer timing experiments (dly3wk = 3-week delay, dly5wk=5 week delay, split = split fertilizer application) at four UAN fertilizer application amounts (N-rate = 0, 56, 112, or 168 kg N $ha^{-1}$ $y^{-1}$).

| **Treatment for sorghum 2019** | **$N_2O$ emissions (kg $N_2O$-N $ha^{-1}$ $yr^{-1}$)** | **$NO_3^-$ leaching (kg $NO_3$-N $ha^{-1}$ $yr^{-1}$)** | **Emissions change vs. noNS at same N-rate (%)** | **Leaching change vs. noNS at same N-rate (%)** | **Emissions change vs. measNS no delay at same N-rate (%)** | **Leaching change vs. measNS no delay at same N-rate (%)** | **Emissions change vs. no fertilizer, measNS (%)** | **Leaching change vs. no fertilizer, measNS (%)** |
|---|---|---|---|---|---|---|---|---|
| 000_noNS_2019 | 7.5 | 16.2 | | | | | | |
| 000_measNS_2019 | 7.4 | 16.2 | -1.4 | -0.0 | | | | |
| 000_maxNS_2019 | 7.2 | 16.2 | -4.2 | -0.1 | | | | |
| 056_noNS_2019 (no delay) | 9.9 | 16.5 | | | | | | |
| 056_measNS_2019 (no delay) | 9.8 | 16.5 | -1.3 | -0.0 | | | +32.2 | +2.2 |
| 056_maxNS_2019 (no delay) | 9.5 | 16.5 | -4.4 | -0.1 | | | +27.9 | +2.2 |
| 056_measNS_dly3wk_2019 | 9.8 | 16.4 | | | -0.3 | -0.5 | +31.8 | +1.7 |
| 056_measNS_dly5wk_2019 | 9.5 | 16.4 | | | -3.2 | -0.7 | +27.9 | +1.6 |
| 056_measNS_split_2019 | 9.4 | 16.5 | | | -4.0 | -0.4 | +26.9 | +1.8 |
| 112_noNS_2019 (no delay) | 12.6 | 16.8 | | | | | | |
| 112_measNS_2019 (no delay) | 12.4 | 16.8 | -1.8 | -0.1 | | | +67.7 | +3.8 |
| 112_maxNS_2019 (no delay) | 11.9 | 16.8 | -5.9 | -0.2 | | | +60.6 | +3.7 |
| 112_measNS_dly3wk_2019 | 12.6 | 16.6 | | | +1.5 | -1.1 | +70.1 | +2.7 |
| 112_measNS_dly5wk_2019 | 12.6 | 16.6 | | | +1.3 | -1.3 | +69.8 | +2.5 |
| 112_measNS_split_2019 | 12.2 | 16.7 | | | -1.6 | -0.6 | +65.0 | +3.2 |
| 168_noNS_2019 (no delay) | 17.0 | 17.0 | | | | | | |
| 168_measNS_2019 (no delay) | 16.7 | 17.0 | -1.8 | -0.1 | | | +125.4 | +5.0 |
| 168_maxNS_2019 (no delay) | 15.3 | 16.9 | -9.8 | -0.4 | | | +107.1 | +4.6 |
| 168_measNS_dly3wk_2019 | 16.9 | 16.7 | | | +1.0 | -1.8 | +127.7 | +3.1 |
| 168_measNS_dly5wk_2019 | 16.6 | 16.5 | | | -0.9 | -2.8 | +123.5 | +2.0 |
| 168_measNS_split_2019 | 16.6 | 16.9 | | | -0.8 | -0.4 | +123.7 | +4.5 |

Table S5. Simulated $N_2O$ emissions, nitrate leaching, and gross nitrification for sorghum in 2018 and sorghum in 2019 comparing fertilizer timing experiments (dly3wk = 3-week delay, dly5wk=5-week delay, split = split fertilizer application) at three UAN fertilizer application amounts (N-rate = 56, 112, or 168 kg N $ha^{-1}$ $y^{-1}$).

| **Treatment for sorghum 2018 or 2019** | **$N_2O$ emissions (kg $N_2O$-N $ha^{-1}$ $yr^{-1}$)** | **$NO_3^-$ leaching (kg $NO_3$-N $ha^{-1}$ $yr^{-1}$)** | **Gross nitrification kg $NO_3$-N $ha^{-1}$ yr-$^{1}$)** | **Emissions change vs. measNS no delay at same N-rate (%)** | **Leaching change vs. measNS no delay at same N-rate (%)** | **Gross nitrification change vs. measNS no delay at same N-rate (%)** |
|---|---|---|---|---|---|---|
| 056_measBNI_2018 (no delay) | 11.8 | 2.7 | 70.9 | | | |
| 112_measBNI_2018 (no delay) | 15.0 | 2.7 | 104.0 | | | |
| 168_measBNI_2018 (no delay) | 18.5 | 2.7 | 141.0 | | | |
| 056_measBNI_dly3wk_2018 | 12.6 | 2.7 | 75.9 | +6.6 | +0.7 | +7.0 |
| 056_measBNI_dly5wk_2018 | 12.2 | 2.7 | 75.1 | +3.5 | +0.5 | +5.9 |
| 056_measBNI_split_2018 | 11.8 | 2.7 | 69.5 | -0.2 | -0.5 | -2.0 |
| 112_measBNI_dly3wk_2018 | 15.8 | 2.7 | 111.2 | +4.9 | -0.4 | +7.0 |
| 112_measBNI_dly5wk_2018 | 15.5 | 2.7 | 110.2 | +3.0 | -0.7 | +5.9 |
| 112_measBNI_split_2018 | 15.2 | 2.7 | 104.9 | +1.2 | -1.1 | +0.8 |
| 168_measBNI_dly3wk_2018 | 19.9 | 2.7 | 150.4 | +7.9 | -0.7 | +6.7 |
| 168_measBNI_dly5wk_2018 | 20.2 | 2.7 | 147.0 | +9.5 | -1.0 | +4.3 |
| 168_measBNI_split_2018 | 18.6 | 2.7 | 142.2 | +0.7 | -0.8 | +0.9 |
| 056_measBNI_2019 (no delay) | 9.8 | 16.5 | 75.5 | | | |
| 112_measBNI_2019 (no delay) | 12.4 | 16.8 | 108.9 | | | |
| 168_measBNI_2019 (no delay) | 16.7 | 17.0 | 147.8 | | | |
| 056_measBNI_dly3wk_2019 | 9.8 | 16.4 | 75.7 | -0.3 | -0.5 | +0.3 |
| 056_measBNI_dly5wk_2019 | 9.5 | 16.4 | 73.5 | -3.2 | -0.7 | -2.7 |
| 056_measBNI_split_2019 | 9.4 | 16.5 | 70.3 | -4.0 | -0.4 | -7.0 |
| 112_measBNI_dly3wk_2019 | 12.6 | 16.6 | 108.1 | +1.5 | -1.1 | -0.7 |
| 112_measBNI_dly5wk_2019 | 12.6 | 16.6 | 104.9 | +1.3 | -1.3 | -3.7 |
| 112_measBNI_split_2019 | 12.2 | 16.7 | 105.9 | -1.6 | -0.6 | -2.7 |
| 168_measBNI_dly3wk_2019 | 16.9 | 16.7 | 138.3 | +1.0 | -1.8 | -6.5 |
| 168_measBNI_dly5wk_2019 | 16.6 | 16.5 | 132.8 | -0.9 | -2.8 | -10.2 |
| 168_measBNI_split_2019 | 16.6 | 16.9 | 145.5 | -0.8 | -0.4 | -1.6 |

Table S6. Simulated $N_2O$ emissions and nitrate leaching for soybean in 2020 that followed sorghum 2019 nitrification suppression experiments (noNS = no nitrification suppression, measNS = measured nitrification suppression, maxNS = maximum nitrification suppression) and fertilizer timing experiments (dly3wk = 3-week delay, dly5wk=5 week delay, split = split fertilizer application) at four UAN fertilizer application amounts (N-rate = 0, 56, 112, or 168 kg N $ha^{-1}$ $y^{-1}$).

| **Treatment for sorghum 2019 that preceded soybean 2020** | **$N_2O$ emissions (kg $N_2O$-N $ha^{-1}$ $yr^{-1}$)** | **$NO_3^-$ leaching (kg $NO_3$-N $ha^{-1}$ $yr^{-1}$)** | **Emissions change vs. noNS at same N-rate (%)** | **Leaching change vs. noNS at same N-rate (%)** | **Emissions change vs. measNS at same N-rate (%)** | **Leaching change vs. measNS at same N-rate (%)** | **Emissions change vs. no fertilizer, measNS (%)** | **Leaching change vs. no fertilizer, measNS (%)** |
|---|---|---|---|---|---|---|---|---|
| 000_noNS_2020 | 5.9 | 6.6 | | | | | | |
| 000_measNS_2020 | 5.9 | 6.6 | +0.0 | -0.2 | | | | |
| 000_maxNS_2020 | 6.0 | 6.6 | +0.2 | -0.6 | | | | |
| 056_noNS_2020 | 6.2 | 7.3 | | | | | | |
| 056_measNS_2020 | 6.2 | 7.2 | +0.0 | -0.2 | | | +4.8 | +9.7 |
| 056_maxNS_2020 | 6.2 | 7.2 | +0.3 | -0.8 | | | +5.0 | +9.1 |
| 056_measNS_dly3wk_2020 | 6.2 | 7.0 | | | -0.8 | -2.6 | +4.0 | +6.8 |
| 056_measNS_dly5wk_2020 | 6.2 | 7.0 | | | -1.1 | -3.4 | +3.7 | +6.0 |
| 056_measNS_split_2020 | 6.2 | 7.1 | | | -1.0 | -2.2 | +3.7 | +7.2 |
| 112_noNS_2020 | 6.5 | 7.8 | | | | | +4.2 | +7.8 |
| 112_measNS_2020 | 6.5 | 7.8 | +0.1 | -0.4 | | | +9.7 | +18.1 |
| 112_maxNS_2020 | 6.6 | 7.7 | +0.6 | -1.2 | | | +10.3 | +17.1 |
| 112_measNS_dly3wk_2020 | 6.5 | 7.4 | | | -1.0 | -4.9 | +8.7 | +12.3 |
| 112_measNS_dly5wk_2020 | 6.5 | 7.4 | | | -0.6 | -5.4 | +9.0 | +11.7 |
| 112_measNS_split_2020 | 6.5 | 7.6 | | | -0.5 | -2.6 | +9.2 | +15.0 |
| 168_noNS_2020 | 6.8 | 8.4 | | | | | +9.4 | +14.8 |
| 168_measNS_2020 | 6.8 | 8.4 | +0.2 | -0.6 | | | +14.6 | +26.7 |
| 168_maxNS_2020 | 7.0 | 8.2 | +2.2 | -2.6 | | | +16.9 | +24.1 |
| 168_measNS_dly3wk_2020 | 6.7 | 8.0 | | | -2.4 | -4.0 | +11.9 | +21.7 |
| 168_measNS_dly5wk_2020 | 6.6 | 7.9 | | | -3.4 | -5.5 | +10.7 | +19.7 |
| 168_measNS_split_2020 | 6.8 | 8.2 | | | +0.1 | -1.8 | +14.7 | +24.4 |

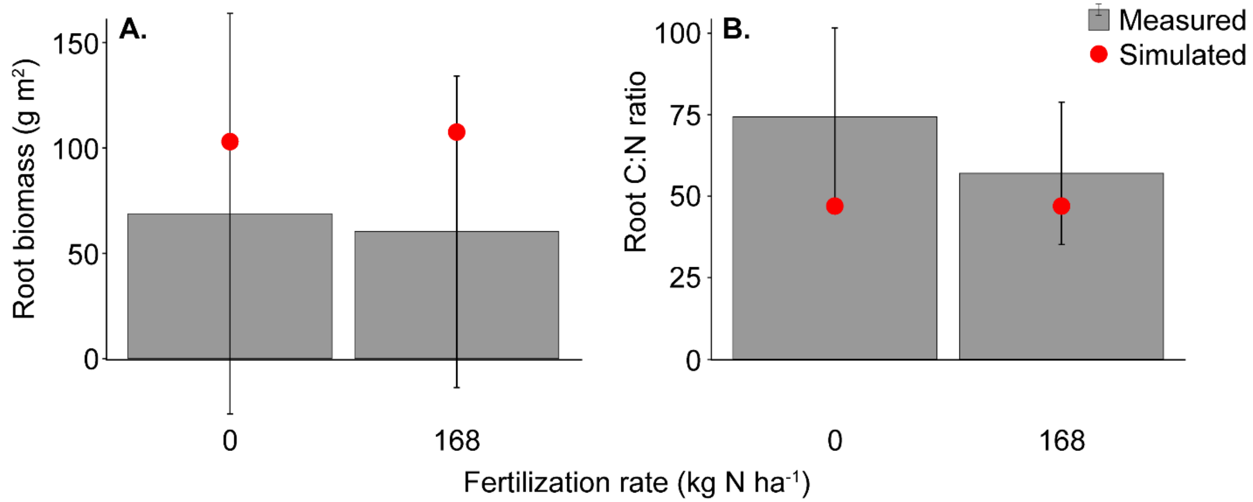


Figure S1. Measured (mean ± 1 standard deviation) and simulated sorghum root biomass (A) and C:N ratio (B) in 2018 across two rates of N fertilization.

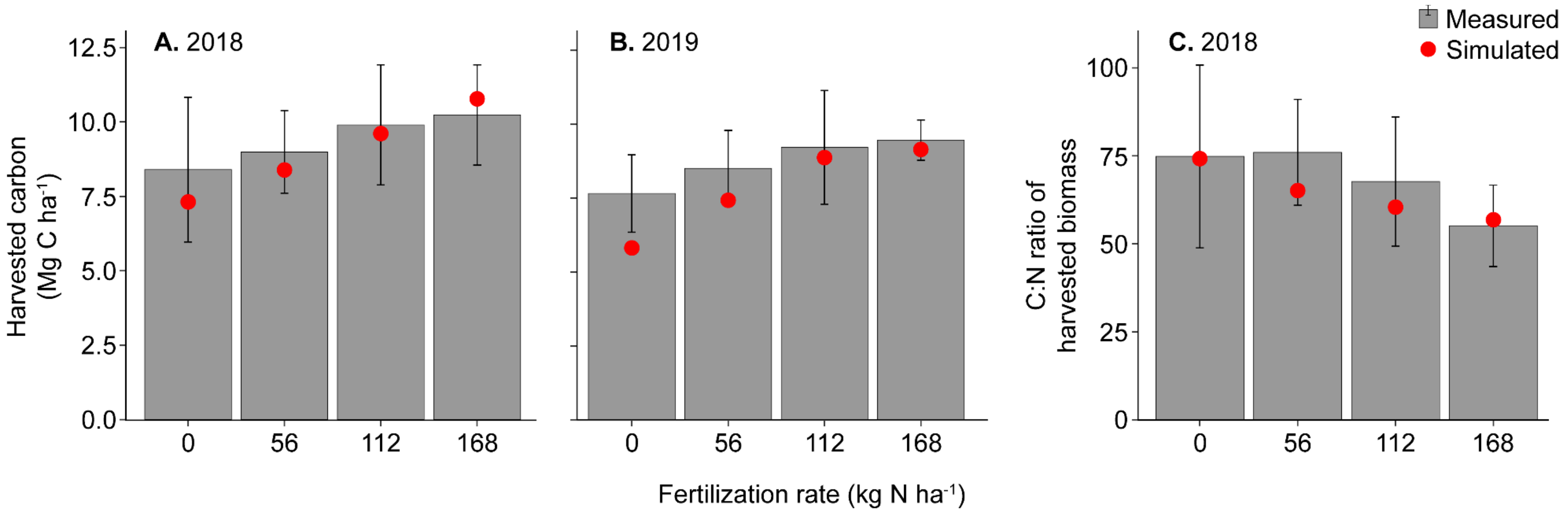


Figure S2. Measured (mean ± 1 standard deviation) and simulated harvested C in sorghum biomass in 2018 (A) and 2019 (B) sorghum trials and C:N ratio of harvested sorghum biomass in 2018 (C) across four rates of N fertilization.

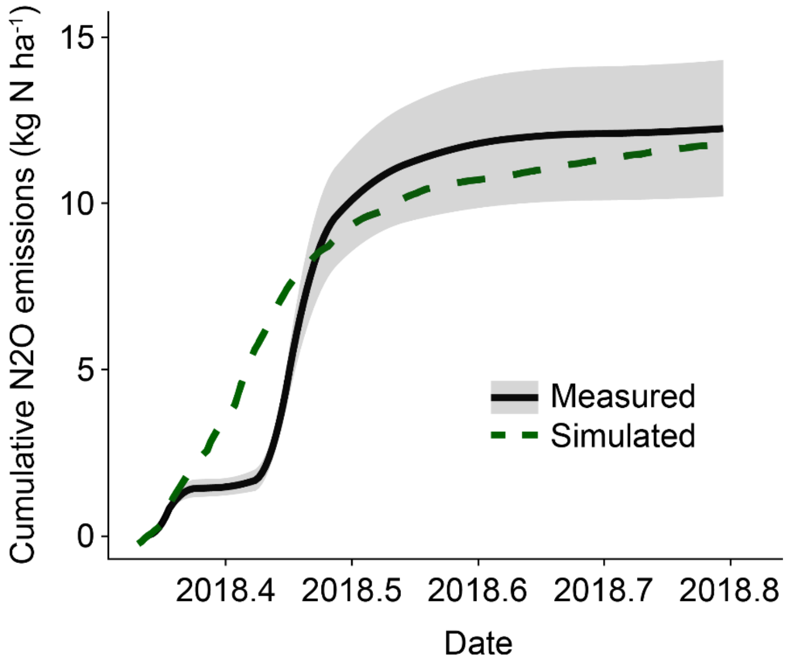


Figure S3. Measured and simulated cumulative $N_2O$ emissions across the 2018 growing season (May 5 – October 18, 2018) in plots fertilized at 112 kg N $ha^{-1}$.

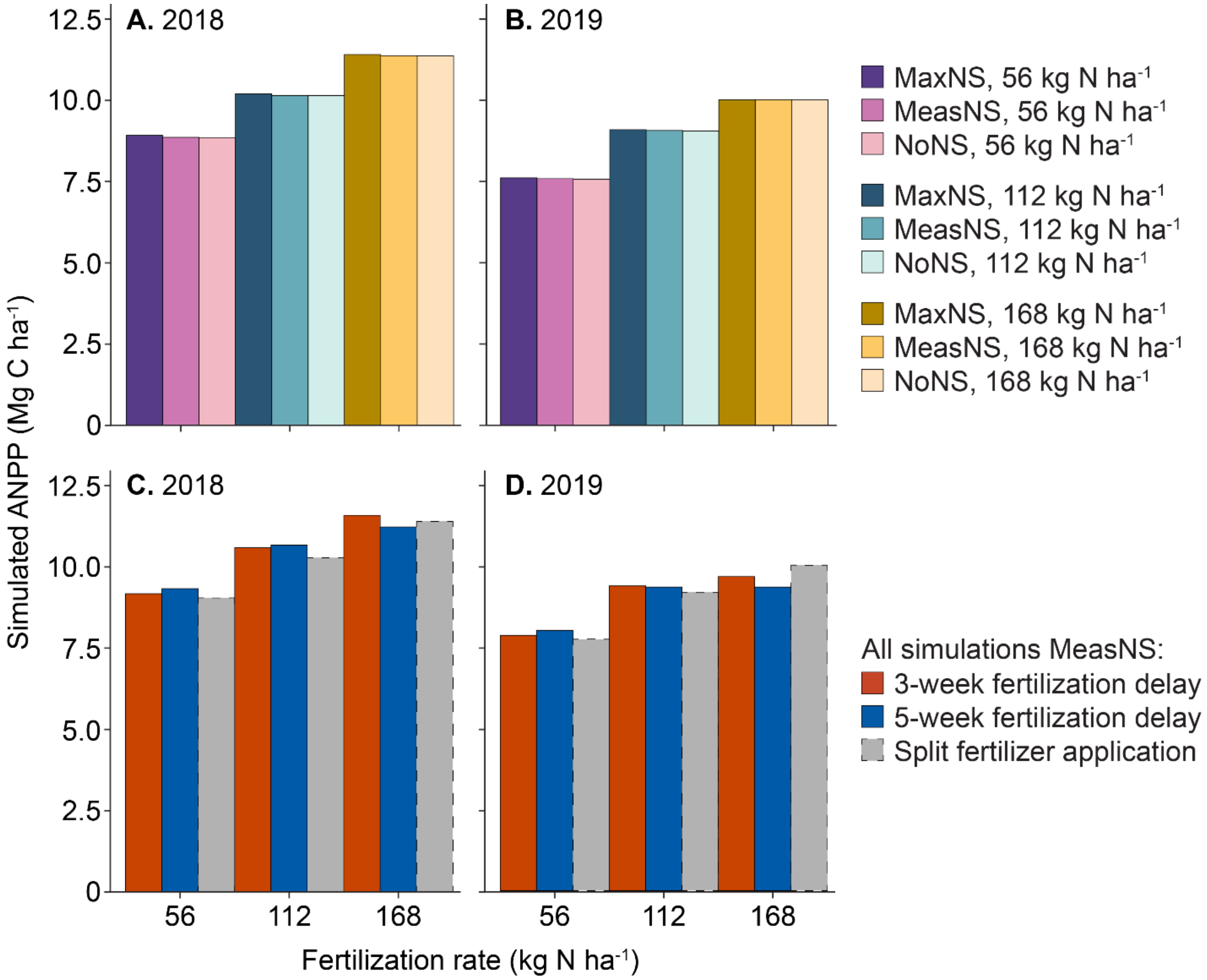


Figure S4. Simulated sorghum aboveground net primary productivity (ANPP, Mg C $ha^{-1}$) in 2018 (A, C) and 2019 (B, D) across three rates of N fertilization for two model experiments. Three levels of simulated nitrification suppression (A, B) include no nitrification suppression (NoNS), measured nitrification suppression (MeasNS), and maximum nitrification suppression (MaxNS). Three fertilization treatments (C, D) include a 3-week and 5-week delay in fertilizer application and a split fertilizer application.

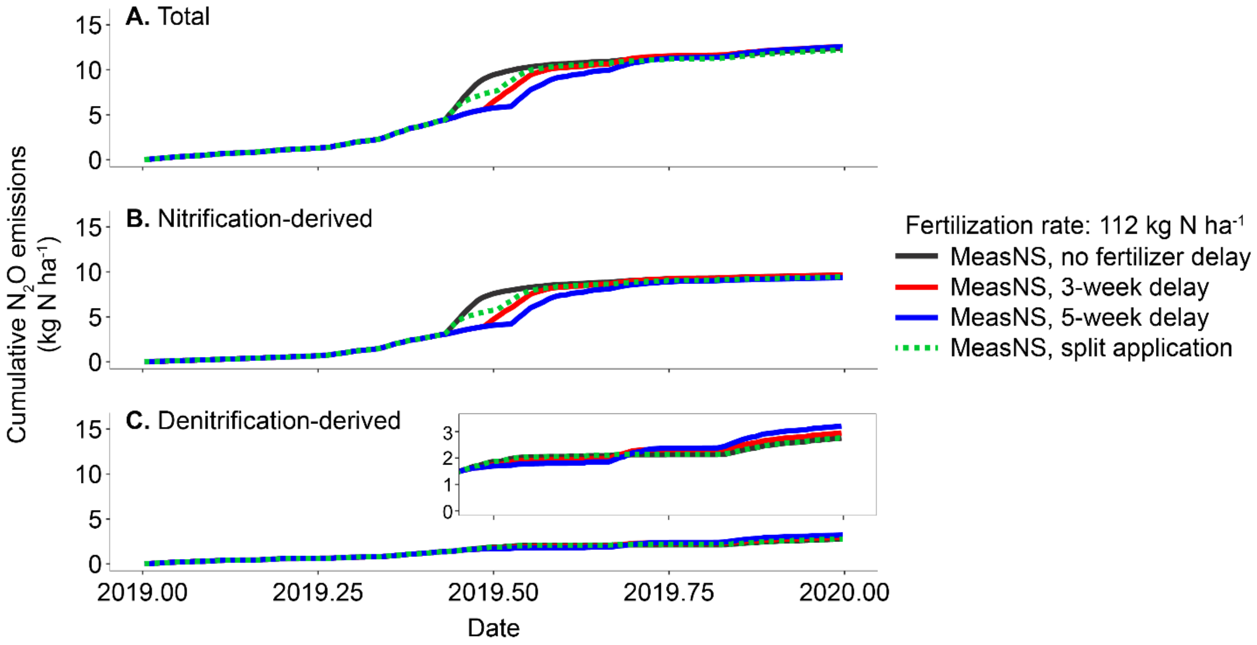


Figure S5. Simulated cumulative total $N_2O$ emissions (A) and $N_2O$ emissions that were derived from nitrification (B) or denitrification (C) across four fertilization application treatments. The inset in panel C shows an expanded vertical axis to illustrate slight differences between fertilizer application treatments. All simulation results are shown at a fertilization rate of 112 kg N $ha^{-1}$ and at measured nitrification suppression (MeasNS).

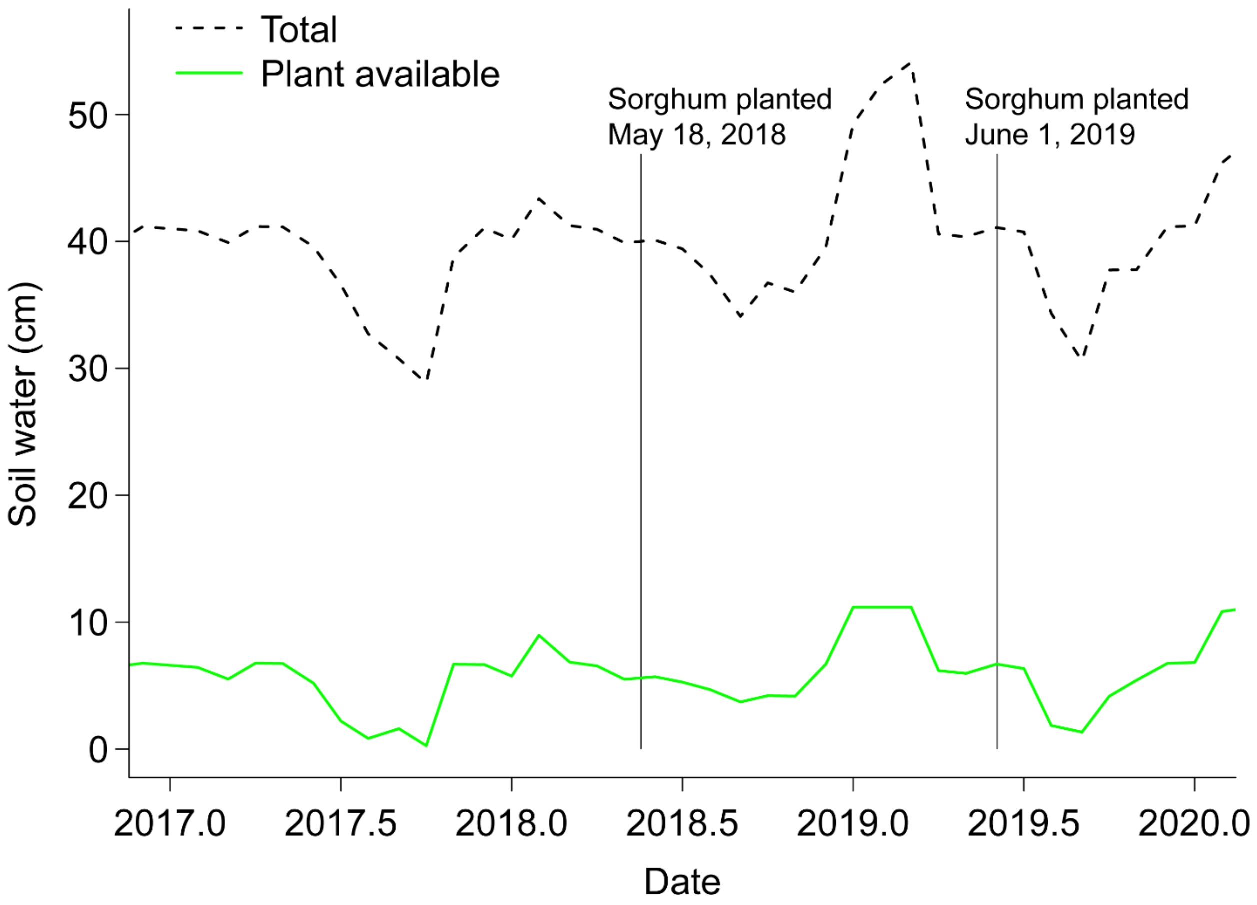


Figure S6. Total soil water that exceeds wilting point and is available to plants for transpiration (green line), and total soil water that exceeds wilting point in the entire soil profile (dashed black line). Vertical lines indicate sorghum planting dates in 2018 and 2019.

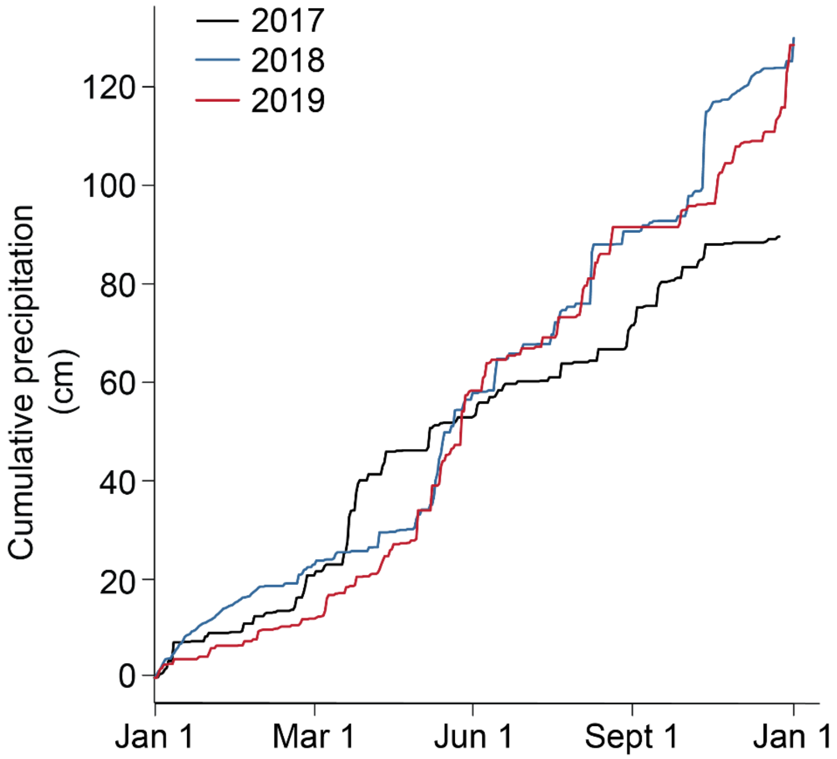


Figure S7. Cumulative precipitation in 2017, 2018, and 2019.
